# A life cycle alteration can correct molting defects in *Caenorhabditis elegans*

**DOI:** 10.1101/2021.11.02.466921

**Authors:** Shaonil Binti, Rosa V. Melinda, Braveen B. Joseph, Phil Edeen, Sam D. Miller, David S. Fay

## Abstract

Molting is a widespread feature in the development of many invertebrates, including nematodes and arthropods. In *Caenorhabditis elegans*, the highly conserved protein kinases NEKL-2/NEK8/9 and NEKL-3/NEK6/7 (NEKLs) promote molting through their involvement in the uptake and intracellular trafficking of epidermal cargos. We found that the relative requirements for NEKL-2 and NEKL-3 differed at different life-cycle stages and under different environmental conditions. Most notably, the transition from the second to the third larval stage (L2→L3 molt) required a higher level of NEKL function than during several other life stages or when animals had experienced starvation at the L1 stage. Specifically, larvae that entered the pre-dauer L2d stage could escape molting defects when transiting to the (non-dauer) L3 stage. Consistent with this, mutations that promote entry into L2d suppressed *nekl*-associated molting defects, whereas mutations that inhibit L2d entry reduced starvation-mediated suppression. We further showed that loss or reduction of NEKL functions led to defects in the transcription of cyclically expressed molting genes, many of which are under the control of systemic steroid hormone regulation. Moreover, the timing and severity of these transcriptional defects correlated closely with the strength of *nekl* alleles and with their stage of arrest. Interestingly, transit through L2d rescued *nekl*-associated expression defects in suppressed worms, providing an example of how life-cycle decisions can impact subsequent developmental events. Given that NEKLs are implicated in the uptake of sterols by the epidermis, we propose that loss of NEKLs leads to a physiological reduction in steroid-hormone signaling and consequent defects in the transcription of genes required for molting.

## Introduction

Developmental plasticity is critical for ensuring the survival of an organism in response to rapidly changing environments (Suzuki & Nijhout, 2008). The dauer larva, a developmentally arrested stage of *C. elegans*, is a classic example of phenotypic plasticity that allows the worm to cope with the unpredictability of nature (Cassada & Russell, 1975; Fielenbach & Antebi, 2008; Golden & Riddle, 1984b). Under permissive conditions after hatching from the egg, *C. elegans* develops through four larval stages (L1, L2, L3, and L4) and eventually becomes an adult (Figure 1A). Conversely, when the conditions are non-optimal (e.g., high temperature, crowding, limited food availability), late-stage L1 larvae sense the harsh environment and initiate a full-body response to form a dauer (also referred to as diapause) larva, an alternative form of L3 that is stress resistant, developmentally quiescent, and designed for survival and dispersion (Cassada & Russell, 1975; Golden & Riddle, 1984b). During the process of forming a dauer, the L1 larva first molts into a pre-dauer L2d larva Figure 1A). The decision to develop into either an L2 or L2d larva happens several hours before the L1→L2(d) molt (Golden & Riddle, 1984b; Schaedel et al., 2012). An L2d larva then has two options—either proceed to form a dauer or, if conditions improve, molt into an L3 larva and resume normal development. This decision is made at the mid-L2d stage, at which point the larva becomes committed to forming a dauer (L3d) or proceeding with L3 development (Figure 1A) (Schaedel et al., 2012). The initial decision to enter L2d from L1 provides a safety net, giving animals the ability to enter dauer (L2d→L3d) should conditions fail to improve, while still allowing animals to resume with normal development (L2d→L3) if the environment becomes permissive. One cost associated with L2d, however, is that that the L2d stage takes several hours longer to complete than L2, thus lengthening the time to become a fertile adult (Golden & Riddle, 1984b; Karp, 2018). Other potential costs or benefits of the L2d decision on subsequent developmental events are not well understood.

**Figure 1.**
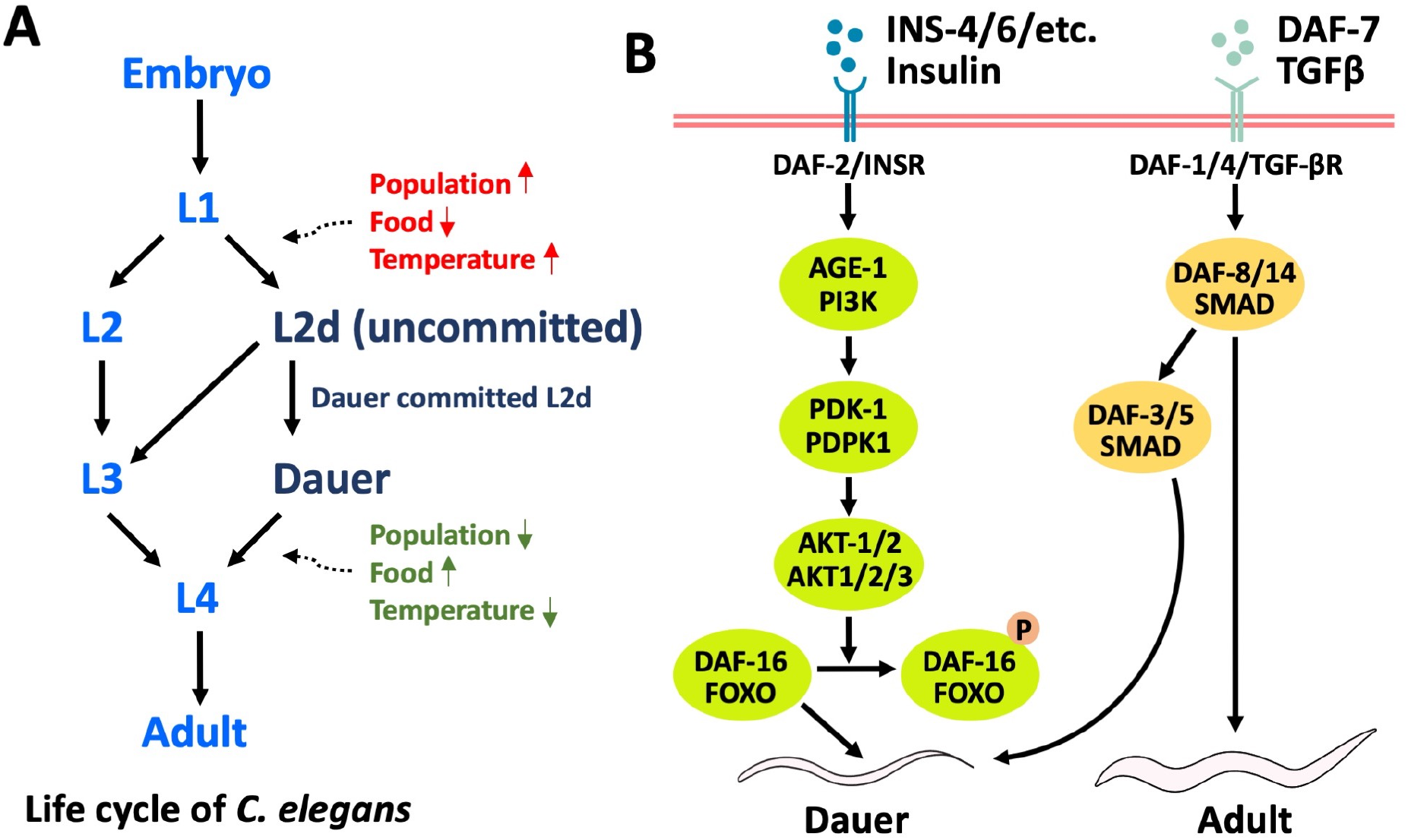
Developmental trajectories of *C. elegans*. (A) Under ideal conditions, an L1 larva develops continuously through four larval stages (L1, L2, L3, L4) to reach adulthood. Under unfavorable conditions (e.g., high temperature, high population density, or limited food availability), an L1 larva molts into an uncommitted L2d pre-dauer larva. If conditions improve, the L2d larva molts into an L3 and resumes reproductive development. Alternatively, if unfavorable conditions persist, the L2d becomes committed to forming a dauer (L3d), which exists in a state of arrested development until conditions improve, at which time it completes L3 programs and molts into an L4. (B) insulin/IGF-1 and TGF-β signaling pathways integrate environmental cues, leading to appropriate developmental decisions. In the presence of food, insulin-like peptides bind to the DAF-2/INSR receptor, which further activates the phospholipid kinase AGE-1/PI3K and the serine/threonine kinases PDK-1, AKT-1, and AKT-2. This signaling cascade leads to phosphorylation and nuclear exclusion of the DAF-16/FOXO transcription factor, which is required for the upregulation of genes that promote dauer. Acting in parallel, the TGF-β pathway is sensitive to changes in population density, food availability, and temperature. Under favorable conditions, TGF-β signaling is initiated by the activation of DAF-1/4/TGF-βR receptors by the DAF-7/TGF-β ligand. This signal is transduced through the activation of DAF-8/SMAD2 and DAF-14/SMAD3, which in turn inhibit the positive regulators of dauer induction, DAF-3/SMAD4 and DAF-5/Ski/Sno.

The endocrine signaling network that regulates the developmental switch between reproductive development and dauer formation has been extensively studied in *C. elegans* (Figure 1B). In well-fed animals, insulin/IGF-1–like peptides are secreted by neuroendocrine cells and bind to DAF-2, the *C. elegans* homolog of the insulin/IGF-1 receptor (Hu, 2007; Kimura et al., 1997). Ligand activation of DAF-2 further transduces the nutrient signal by activating downstream kinases that eventually phosphorylate and sequester the DAF-16/FOXO transcription factor in the cytoplasm (Lee et al., 2001; K. Lin, 1997; Kui Lin et al., 2001; Ogg et al., 1997). Reduced activity of the insulin/IGF-1 signaling pathway due to nutrient stress or mutations in upstream genes allows DAF-16 to enter the nucleus and promote the expression of target genes necessary for dauer transformation and arrest. Acting in a semi-parallel fashion to insulin/IGF-1, the TGF-β signaling pathway is responsive to environmental cues such as dauer pheromone, which is an indicator of population density (Golden & Riddle, 1984b; Ren et al., 1996; Riddle & Albert, 1997). Under satisfactory conditions, secreted DAF-7/TGF-β binds to the TGF-β receptors DAF-1 and DAF-4, leading to the inhibition of downstream SMAD proteins DAF-3/SMAD4 and DAF-5/Ski/Sno, which, like DAF-16, promote the dauer program (Fielenbach & Antebi, 2008; Gumienny & Savage-Dunn, 2013). The presence of dauer pheromone in the environment inhibits DAF-7 expression, thus allowing DAF-3 and DAF-5 to favor dauer formation. (Fielenbach & Antebi, 2008; Ren et al., 1996). Acting together, these two pathways integrate diverse environmental cues, leading to an appropriate developmental choice.

Interestingly, worms do retain some memory of prior developmental choices. When compared with an adult that developed continuously, a post-dauer adult exhibits the differential expression of many genes including those associated with reproduction, transcriptional regulation, and metabolism (Hall et al., 2010). Dauer formation also suppresses abnormal germ cell proliferation in adults, suggesting that progression through dauer can alter phenotypes at subsequent life stages (Mao et al., 2020). Correspondingly, passage through dauer can correct spatial and temporal defects in cell fate patterning, indicating that dauer may induce compensatory mechanisms to maintain developmental robustness (Abrahante et al., 2003; Battu et al., 2003; Baugh & Hu, 2020; Ferguson & Horvitz, 1985; Ilbay & Ambros, 2019).

In this study, we report the surprising finding that entry into uncommitted L2d can compensate for a defect in the *C. elegans* molting cycle. Molting is a relatively understudied but crucial developmental process that allows *C. elegans* to grow and progress through larval stages into an adult (Lažetić & Fay, 2017b). During each molt, the old cuticle detaches from the underlying epidermis, while a new cuticle is synthesized underneath. Molting is orchestrated in large part by nuclear hormone receptor signaling, which controls the expression of thousands of genes that oscillate with the molting cycle (Antebi, 2015; Frand et al., 2005; Hendriks et al., 2014; Meeuse et al., 2020; Turek & Bringmann, 2014). One key molecule in this process is cholesterol obtained from the environment, which is required for the synthesis of steroid hormone ligands that activate molting-associated nuclear hormone receptors such as NHR-23/RORC and NHR-25/NR5A1/2 (Asahina et al., 2000; Kostrouchova et al., 2001; Kurzchalia & Ward, 2003; Lažetić & Fay, 2017b; Monsalve & Frand, 2012). Consistent with this, *C. elegans* requires dietary cholesterol for progression through all four larval stages, and cholesterol deprivation impairs growth and other physiological processes including molting (Shim et al., 2002; Yochem et al., 1999). The importance of steroid hormone signaling is also underscored by the observation that the epidermally expressed low-density lipoprotein (LDL)-like receptor LRP-1/Megalin is required for molting and is thought to mediate the uptake of sterols by the epidermis.

We previously showed that the NIMA-related kinases NEKL-2/NEK8/9 and NEKL-3/NEK6/7 (NEKLs)—along with their conserved ankyrin repeat binding partners, MLT-2/ANKS6, MLT-3/ANKS3, and MLT-4/INVS (MLTs)—promote molting through their roles in regulating the trafficking of epidermal cargos. In larvae harboring mutations in *nekl* and *mlt* genes, the old cuticle fails to detach fully during molting and worms undergo larval arrest (Lažetić & Fay, 2017a; Yochem et al., 2015). Within the epidermis, depletion of NEKLs leads to impaired clathrin-mediated endocytosis and reduced internalization of LRP-1 (Joseph et al., 2020). Here we show that downregulation of NEKL*s* results in the delayed and/or reduced expression of hormonally regulated molting-specific genes, thereby establishing a link between NEKL trafficking functions and the expression of genes required for molting. Notably, this defect can in some circumstances be overcome by entry into L2d, which, together with other data, indicates clear differences in the requirements for NEKLs at different molts along with demonstrating the impact of life-cycle decisions on downstream phenotypic outcomes.

## Materials and Methods

### Strains and maintenance

*C. elegans* strains were cultured on nematode growth medium (NGM) spotted with *E. coli* OP50 and maintained at 20°C, unless noted otherwise. Synchronous populations of L1 larvae were obtained by bleaching gravid adults (Figures 7A, 8, and S4–S7; Porta-de-la-Riva et al., 2012). After the bleach treatment, eggs were washed five times and allowed to hatch in M9 buffer overnight at room temperature. Hatched worms remain arrested as L1 larvae in the M9 buffer until they are transferred to a plate containing food. All strains used in this study are listed in File S1.

### CRISPR/Cas9-generated loss-of-function mutation in *daf-16*

The *fd335* allele of *daf-16* was created using the *dpy-10* co-CRISPR approach (Arribere et al., 2014; Kim et al., 2014; Paix et al., 2014, 2015). The *fd335* allele contains two premature stop codons at (K248Stop and G250Stop) in DAF-16a, along with an introduced restriction site.

Primer sequences for the sgRNA and repair template are given below. Capital letters represent the altered or inserted nucleotides.

sgRNA for *daf-16(fd335)*: gtacgccgtggattccttcc

Repair Template:

attcagaatgaaggagccggaaagagctcgtggtgggttattaatccagaAgcTTagccaTgaaggaatcacggcgtacacgtgaacgatccaatactattgagacgactacaaaggtaagaga

*nekl-2(fd81)*; *daf-16(fd335)*; *nekl-3(gk894345)*; *fdEx286* worms were allowed to starve and were subsequently examined for the presence of dauers. We failed to detect (GFP^+^) dauer worms based on morphology. In addition, we failed to detect any surviving worms following treatment with 1% SDS, further indicating that *fd335* blocks dauer formation.

### Image acquisition

For the auxin treatment experiments in Figure S3, fluorescence images were obtained using an Olympus IX83 P2ZF inverted microscope. Z-stacks were captured with a step size of 0.2 µm, using a 100 × silicon oil immersion objective (1.35 N.A.). Images in Figures S5 and S6 were taken with an Olympus MVX10 MacroView microscope using a 2 ×objective (0.5 N.A.). The rest of the DIC images and fluorescence images were obtained using a Nikon Eclipse epifluorescence microscope using 10 × (0.25 N.A.) and 40 (0.75 N.A.) objectives. Images were acquired using Olympus cellSens software version 3.1. Before imaging, animals were mounted on 3% agarose pads and immobilized using 10 mM levamisole.

### Image analysis

All the image processing and analyses were performed using FIJI software (Schindelin et al., 2012). For the auxin treatment experiments, the fluorescence images were first background subtracted, and then image thresholding was performed using the Li algorithm (Li & Tam, 1998). A representative region of interest in hyp7 was selected that covered most of hyp7 in the frame. Then the percentage of positive pixels was quantified for the selected region of interest.

### Auxin-mediated degradation studies

The protocol for the auxin treatment experiments was performed as described in our previous paper (Joseph et al., 2020). To generate the graphs on the left in Figure 6C–E, development was synchronized by bleaching (Porta-de-la-Riva et al., 2012), except for the synchronization of L3 and L4 stages of *nekl-3::aid* genotype which was done manually by characterizing vulval and gonad development. Animals were treated with auxin during the early period of larval development at L1, L2, L3, and L4 stages. Identification of larval stages was performed by following gonad morphology under the dissecting scope. To generate the graphs on the right in Figure 6C–E, starved worms were gently washed off the plate and treated with 1% SDS to isolate dauer larvae (Karp, 2018). Isolated dauers were transferred to an unseeded plate (i.e., one without a bacterial lawn) and treated with auxin for 20 hours. After the auxin treatment, dauers were transferred back to a seeded plate (without auxin) to allow them to grow.

For the experiments in Figure S3, all three strains (wild type, *nekl-3::aid*, and *daf-2*; *nekl-3::aid*) were cultured at 22°C until *daf-2*; *nekl-3::aid* mutants formed dauers. Dauer larvae were identified based on morphological characteristics and transferred to a separate plate. Strains were then incubated at 16°C until *daf-2*; *nekl-3::aid* dauers molted into L4s. From these plates, L4 larvae were selected and transferred to separate plates where they continued to develop into adults. Day-1 adults of wild type, *nekl-3::aid*, and *daf-2*; *nekl-3::aid* genotypes were treated with auxin for 20 hours before imaging. For each experiment, at least 30 worms were imaged for the expression of LRP-1::GFP in the hyp7 epidermis.

### Dauer pheromone experiments

For dauer pheromone experiments, NGM plates were made without peptone. Synthetic ascarosides (ascr#2, ascr#3, and ascr#5), which were kindly provided by the Schroeder lab (Butcher et al., 2008), were added to the liquid medium before it was poured into the plates, so that the final concentration of each ascaroside was 3 μM (Ilbay & Ambros, 2019). Poured plates were spotted with heat-killed *E. coli* OP50. To prepare the bacteria, 5 mL of an overnight culture of *Escherichia coli* OP50 was pelleted, washed twice with sterile water, and dissolved in a tube containing 1 mL of sterile water. The tube was then placed in a water bath set at 95°C for 30 min and vortexed vigorously every 5 min during this incubation (Hollister et al., 2013). Six gravid adults were placed onto an unseeded plate to clear off excess live *E. coli*. They were then transferred to dauer pheromone–containing plates and allowed to lay eggs for 4 hours before being removed. Worms were grown on dauer pheromone plates until *nekl-2*; *nekl-3* (GFP^+^) worms formed dauers. Larvae were then washed off the plates with M9 and transferred to fresh NGM plates spotted with live *E. coli. nekl-2*; *nekl-3* (GFP^−^) worms were scored for molting defects after ∼48 hours.

### Pharyngeal pumping assay

Gravid adults were treated with bleach to generate synchronized populations of L1 larvae as described above. Pharyngeal pumping was scored for >50 synchronized larvae every hour at 20°C after release from L1 arrest. An Olympus MVX10 MacroView microscope was used with a 2 × objective to observe and score pharyngeal pumping of worms. Additional information is in File S2.

### Transcriptional analysis of molting genes

Synchronized L1 larvae (obtained as described above) were cultured at 20°C after release from L1 arrest. Ten worms were scored for the expression of reporter genes every hour using the Olympus MVX10 MacroView microscope with a 2 × objective. Worms were scored as positive or negative for marker expression, leading to the determination of the percentage of worms expressing (% expression) used in Figures 8 and S4. Reduced/inconsistent expression was still scored as positive for the marker expression.

### Statistical Analysis

GraphPad Prism 9 software was used to perform all statistical analyses and to generate bar graphs and line plots.

## Results

### Starvation as a suppressor of *nekl-2*; *nekl-3* molting defects

We previously identified weak loss-of-function alleles of *nekl-2*(*fd81*) and *nekl-3*(*gk894345*) that do not exhibit molting defects on their own but, when combined, exhibit synthetic lethality with ∼98.5% of worms arresting with molting defects (Figure 2A) (Lažetić & Fay, 2017a). Most typically, *nekl-2(fd81*); *nekl-3(gk894345)* (hereafter *nekl-2*; *nekl-3*) mutants transit the L1→L2 molt but then arrest at the L2→L3 molting stage. Arrested worms exhibit a partial cuticle-shedding defect (the “corset” phenotype), in which the old cuticle fails to detach from the midbody region (Yochem et al., 2015). *nekl-2*; *nekl-3* double mutants are maintained by the presence of an extrachromosomal array (*fdEx286*) that expresses wild-type *nekl-3* along with a broadly expressed GFP reporter (Figure 2A).

**Figure 2.**
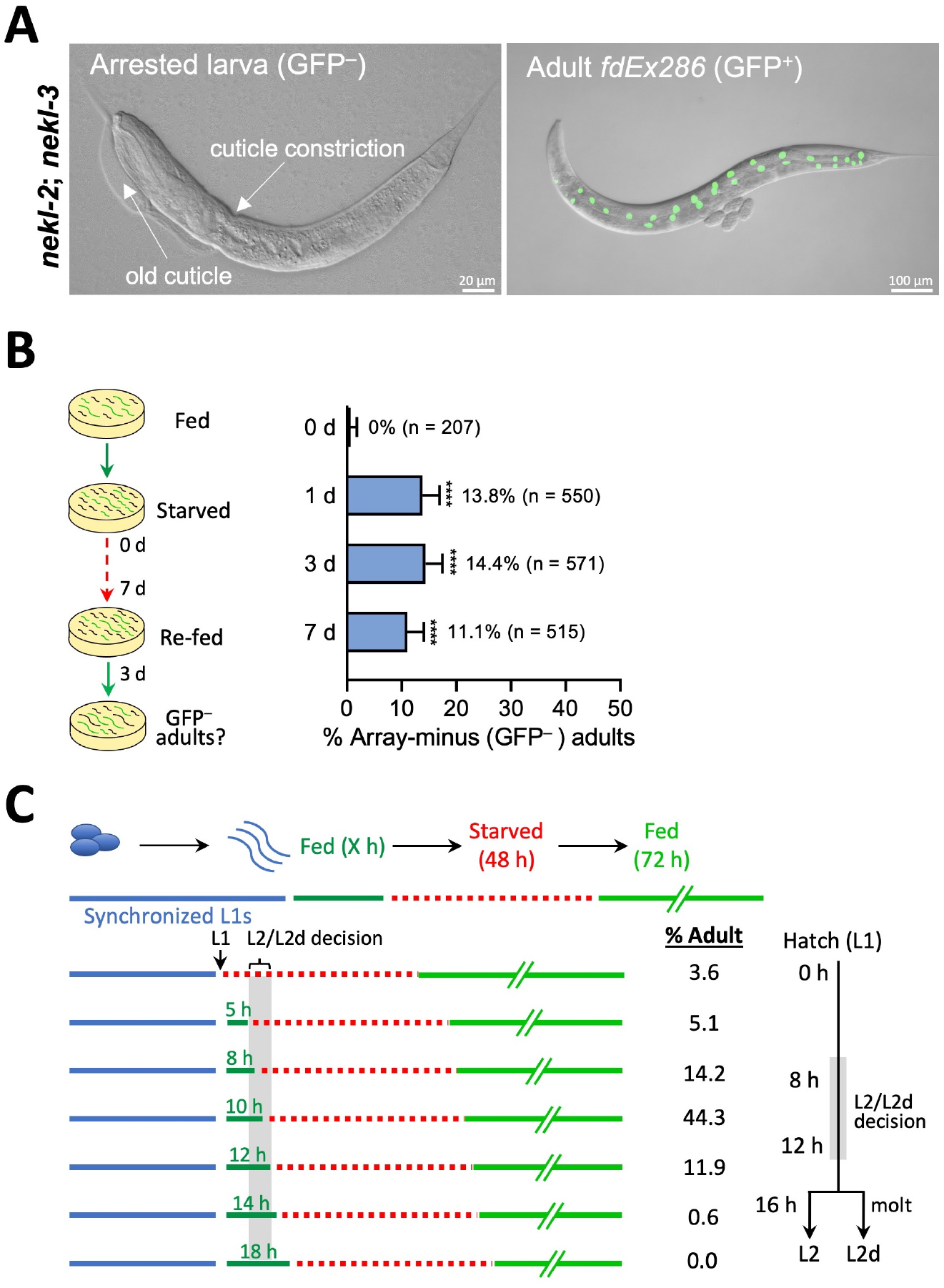
Starvation-induced suppression of molting defects in *nekl-2; nekl-3* mutants. (A) Representative images (DIC merged with green fluorescence) of *nekl-2*; *nekl-3* double mutants without (left) and with (right) the rescuing extrachromosomal array (*fdEx286*), which encodes wild-type NEKL-3 and the SUR-5::GFP reporter. White arrows indicate the position of the constriction where the old cuticle failed to detach from the epidermis (double cuticle) and a portion of the old anterior cuticle that is still partially attached to the worm. (B) Mixed-stage populations of *nekl-2*; *nekl-3* double mutants were starved for 0, 1, 3, or 7 days. The number of viable array-minus (GFP^−^) adults and array-positive (GFP^+^) adults were counted after worms were transferred to a plate containing food and allowed to grow for 3 days. Bar graph represents the percentage of array-minus (GFP^−^) adults observed in the total adult population (GFP^+^+ GFP^−^). The percentage of *nekl-2*; *nekl-3* (GFP^−^) viable adults after 1, 3, and 7 days of starvation was statistically significant as compared with 0 days (^****^p < 0.0001 as determined by Fisher’s exact test). The total number of adult worms is shown in parentheses after each percentage. (C) Synchronized *nekl-2*; *nekl-3* L1 larvae were fed for variable lengths of time (X hours) before being starved for 48 hours and then returned to plates containing food for 72 hours. The percentage of viable GFP^−^ adults, [GFP^−^ adults ÷(GFP^−^ adults + GFP^−^ arrested larvae) 100], was then scored. A corresponding timeline showing the period of the L2→L2d decision is shown to the right.

In our studies, we observed that starved plates containing *nekl-2*; *nekl-3* larvae produced elevated levels of array-minus (GFP^−^) adults following transfer to plates containing bacterial food. To quantify this phenomenon and to determine if starvation length is a factor in suppression, we first assayed non-synchronized worm populations from plates that had been starved for 0, 1, 3, or 7 days prior to food reintroduction (Figure 2B). Notably, starvation of *nekl-2*; *nekl-3* double mutants for 1, 3, or 7 days led to a highly significant increase in the percentage of *nekl-2*; *nekl-3* (GFP^−^) animals that reached adulthood as compared with the absence of starvation (Figure 2B). Furthermore, we observed no significant differences in *nekl-2*; *nekl-3* suppression between plates that had been starved for 1, 3, or 7 days, indicating that the suppressive effects of starvation occur within 24 hours (Figure 2B).

### Environmentally induced L2d can alleviate molting defects in *nekl-2*; *nekl-3* mutants

To better define the crucial developmental window for starvation-mediated suppression, we synchronized newly hatched *nekl-2*; *nekl-3* L1 larvae and then transferred them to plates containing food for varying amounts of time. Larvae were then washed and moved to plates lacking food for 48 hours to induce a starvation response. Finally, larvae were returned to plates containing food for 72 hours to allow for the growth of suppressed worms to adulthood (Figure 2C). Interestingly, we found that induction of a starvation response, after the larvae were first grown for 10 hours on food, led to the highest levels of suppression, whereas starvation after 8 or 12 hours on fed plates led to moderate levels of suppression (Figure 2C). In contrast, starvation after growth on food for 14 or 18 hours failed to induce any suppression, and starvation directly after hatching (0 hours) or after 5 hours on food led to only low levels of suppression (Figure 2C). Notably, the optimal period for inducing suppression corresponded with the known time point at which L1 larvae become committed to either the L2 or L2d developmental programs (Golden & Riddle, 1984b). Moreover, larvae that experienced starvation after the L2→L2d decision point (e.g., 14 and 18 hours) were unable to enter L2d, consistent with the absence of suppression. Taken together, our findings suggest that progression through L2d, but not starvation per se, can alleviate molting defects in *nekl-2*; *nekl-3* worms. Moreover, the ability of the L1-specific starvation regimen to reduce *nekl-2*; *nekl-3* molting defects indicates that uncommitted L2d, but not committed L2d or dauer entry, is sufficient for suppression.

To further test if the L2d developmental trajectory can suppress *nekl* molting defects, we used a mixture of ascarosides (collectively known as dauer pheromone) that induce a high percentage of L2d and dauers in wild type (Butcher et al., 2008). We found that ∼ 17% of the *nekl-2*; *nekl-3* double mutants were able to reach adulthood when briefly exposed to ascarosides (Figure 3A), consistent with the model that *nekl-2*; *nekl-3* mutants can be suppressed by transit through L2d. Furthermore, two findings confirmed that the conditions used here were effective at triggering progression through L2d/dauer. (1) Treatment of worms carrying a validated marker for dauer and pre-dauer stages, *P*_*ets-10*_*::GFP*, resulted in expression patterns expected for uncommitted L2d, committed L2d, and dauer stages (Figures 3B and S1; Shih et al., 2019). (2) *nekl-2; nekl-3* double mutants carrying the *nekl-3*^*+*^; *GFP*^*+*^ rescuing array readily formed dauers in response to ascarosides as well as starvation. We note, however, that (GFP^−^) *nekl-2*; *nekl-3* double mutants uniformly failed to form dauers when subjected to starvation (n > 1000), indicating that *nekl-2*; *nekl-3* double mutants are at least partially defective in the L2d→dauer transition. Curiously, when treated with ascarosides, ∼70% of *nekl-2*; *nekl-3* mutants (n = 228) were able to form dauers, suggesting that the strength or method of the dauer induction can impact the ability of *nekl-2*; *nekl-3* mutants to execute the L2d→dauer transition.

**Figure 3.**
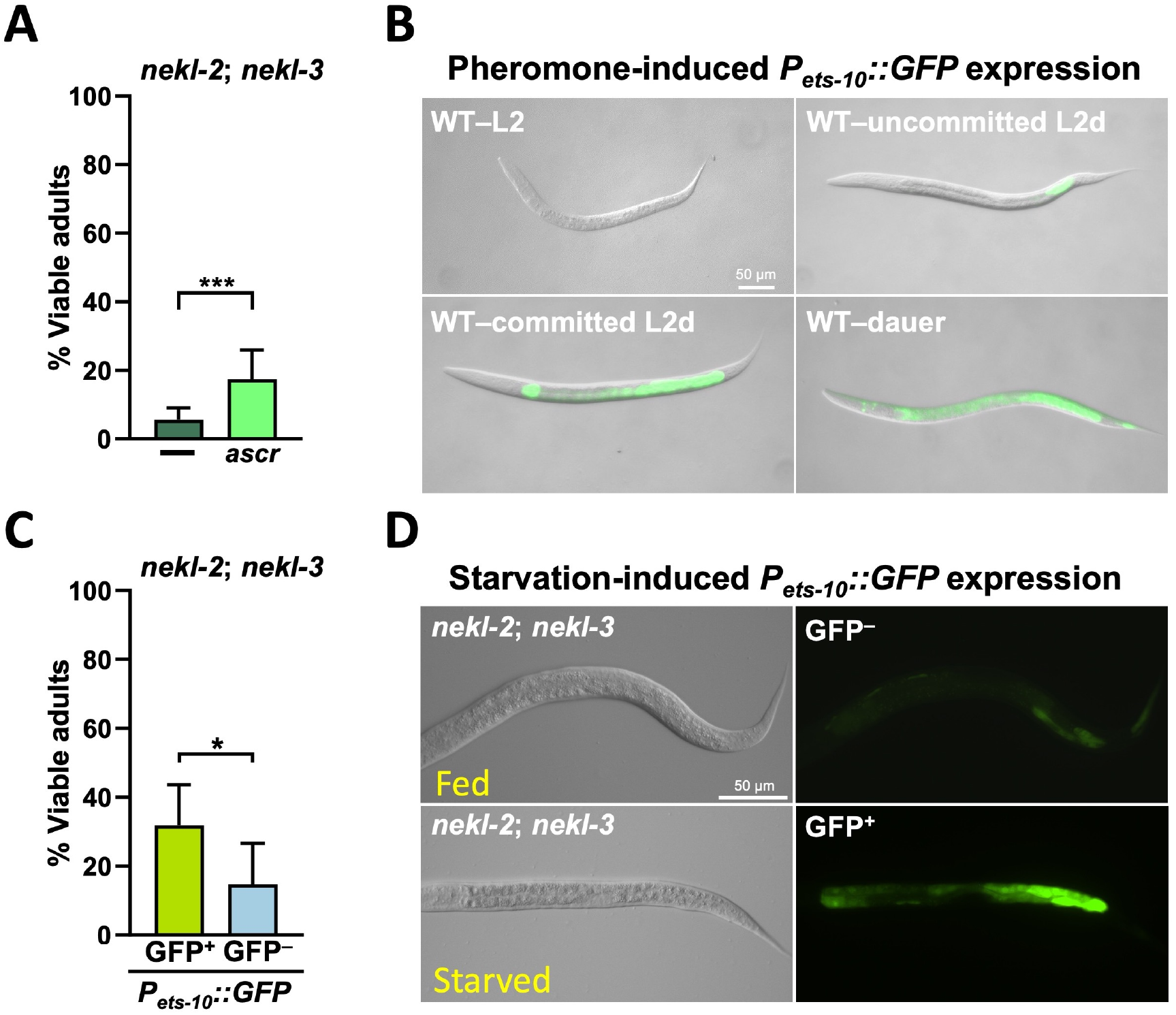
L2d induced by dauer pheromone or starvation suppresses molting defects. (A) Graph showing the percentage of viable *nekl-2*; *nekl-3* adults in the absence and presence of dauer-promoting ascarosides (ascr). (B) Worms carrying an integrated *P*_*ets-10*_*::GFP* reporter grown on ascaroside-containing plates exhibit a differential pattern of expression during dauer formation. Expression of *P*_*ets-10*_*::GFP* was limited to the posterior intestine of uncommitted L2d larvae but then spread throughout the intestine as L2d larvae became committed to forming dauers. (C) Late-stage L1 and early-stage L2 *nekl-2*; *nekl-3* larvae were starved for 1 day, transferred to food plates, and scored/sorted for expression of *P*_*ets-10*_*::GFP* under a high-power dissecting microscope before being transferred to plates with bacteria. Worms were then assessed several days later for progression to adulthood. (D) Representative images of *nekl-2*; *nekl-3* mutants (roughly staged as late L1 to L2) expressing *P*_*ets-10*_*::GFP* under well-fed (upper) and starved (lower) conditions. Error bars represent 95% confidence intervals. All statistical tests were performed using Fisher’s exact test; ^*^p **<** 0.05, ^****^p < 0.001.

To further strengthen the link between L2d and the suppression of molting, we crossed the *P*_*ets-10*_*::GFP* marker into *nekl-2*; *nekl-3* mutants. Late-stage L1 or early-stage L2 *nekl-2*; *nekl-*3 larvae were identified from a population that had experienced starvation for 1 day. We next sorted larvae based on whether they expressed *P*_*ets-10*_*::GFP* in the uncommitted-L2d pattern and categorized the larvae as GFP^+^ or GFP^−^ before being moved to plates spotted with live *E. coli* (Figure 3D). Notably, starved GFP^+^larvae were more likely to mature into adults as compared with their GFP^−^ siblings (Figure 3C). This difference was significant even though it is likely that some of the GFP^−^ worms had entered L2d but that the marker was too dim to be detected by our methods. Taken together, our findings indicate that L2d larvae have a higher tendency to overcome *nekl* molting defects and reach adulthood as compared with L2 larvae.

### Dauer-constitutive mutations alleviate *nekl* molting defects

We next determined if the dauer-inducing signals of dauer-constitutive (Daf-c) mutations could also suppress molting defects in *nekl-2*; *nekl-3* mutants. Downregulation of DAF-2/IGF1R and DAF-7/TGF-β promote dauer entry by de-repressing downstream dauer-promoting transcription factors (Figure 1B). As such, loss-of-function mutations in *daf-2* and *daf-7* lead to worms forming dauers even in the presence of ample food (Gems et al., 1998; Karp, 2018; Riddle & Albert, 1997). Because dauer formation is a temperature-sensitive process (Golden & Riddle, 1984a), we scored for suppression of molting defects by *daf-2* at two different temperatures, 16°C and 20°C. Loss of function of *daf-2* resulted in ∼30% of *nekl-2*; *nekl-3* mutants reaching adulthood at 20°C, whereas no suppression was observed at 16°C (Figure 4A–B). Likewise, we observed robust suppression (∼60%) of molting defects by *daf-7* at 20°C (Figure 4B–C). We note that because *nekl-2(fd81)* is itself a temperature-sensitive allele, we were unable to perform suppression assays at 25°C (also see below). Furthermore, consistent with our prior observations under starvation conditions, the large majority of *nekl-2*; *daf-2*; *nekl-3* mutants failed to form mature dauers, indicating a requirement for NEKLs in the L2d→dauer transition. Together, these results suggest that triple mutants entered L2d at 20°C but then resumed reproductive development and that progression through L2d→L3 was sufficient to alleviate molting defects.

**Figure 4.**
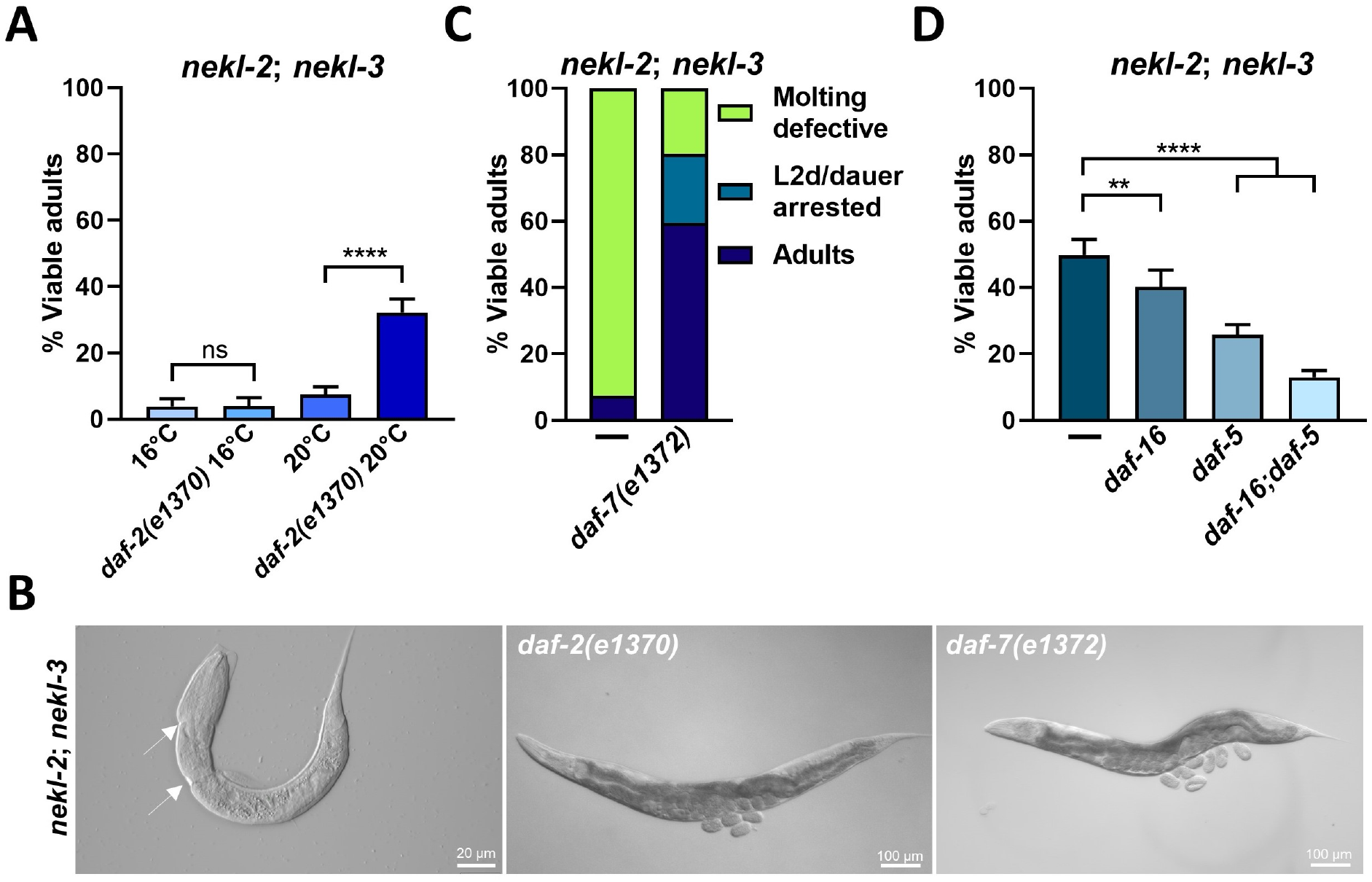
Genetic analysis is consistent with L2d-mediated suppression of molting defects. (A) Bar graph showing the percentage of viable adults when *nekl-2*; *nekl-3* or *nekl-2*; *daf-2*; *nekl-3* mutants were grown at 16°C or 20°C. (B) DIC images of the mutants assessed in panel A at 20°C. White arrows show region(s) of constriction where the old cuticle has failed to detach from the epidermis. (C) Graph showing viable adults (%) among *nekl-2*; *nekl-3* animals in the absence or presence of the *daf-7* mutation. As strains containing *daf-7(e1372)* form dauers at 16°C, we scored suppression at 20°C only. (D) *nekl-2*; *nekl-3* mutants in the presence of Daf-d mutations *daf-5* and *daf-16*, were allowed to starve for three days. Then strains were chunked to new plates and the percentage of viable adults was scored after ∼ 48 hours. Error bars represent 95% confidence intervals. Statistical significance was determined using Fisher’s exact test; ^**^p < 0.01, ^****^p < 0.0001. ns, not significant.

Given our previous data showing that starvation-induced L2d can suppress molting defects in *nekl-2*; *nekl-3* mutants, we were curious to determine whether mutations that prevent dauer formation would also prevent suppression by starvation. To test this, we assayed *nekl-2*; *nekl-3* suppression by starvation in *daf-16, daf-5*, and *daf-16*; *daf-5* dauer-defective (Daf-d) backgrounds. Interestingly, although the CRISPR-generated allele *daf-16(fd335)* prevented the induction of dauer (see Materials and Methods), it resulted in only a minor reduction in suppression by starvation in the *nekl-2*; *nekl-3* background (Figure 4D). Consistent with this, suppression of *nekl-2*; *nekl-3* by the Daf-c allele *daf-2(e1370)* was not significantly reversed by the Daf-d *daf-16(fd335)* mutation (Figure S2). In the case of *daf-5*, we observed a reduction in starvation-induced suppression by ∼ 2-fold, whereas the *daf-16*; *daf-5* combination reduced suppression by ∼ 5-fold. Thus, whereas Daf-d mutations reduced the effectiveness of starvation-mediated suppression to varying extents, they did not entirely prevent it. This may be because these mutations, while fully preventing dauer formation, do not completely prevent entry into uncommitted-L2d. Collectively, our genetic data support a model of *nekl-2*; *nekl-3* suppression that relies on entry into uncommitted L2d and suggest that Daf-d mutations may retain some ability to enter L2d or a partial L2d-like state.

### The L2d→L3 developmental trajectory is not sufficient to suppress strong loss-of-function *nekl* mutations

We next determined whether starvation can suppress developmental arrest in other *nekl* and *mlt* alleles that exhibit molting defects at either the L1→L2 or L2→L3 transitions. We did not observe any suppression of L1→L2 molting defects in strong or null alleles of *nekl-2, nekl-3*, and *mlt-3* (Figure 5A). This result was expected, as mutant larvae become arrested prior to entering L2d. Interestingly, we also observed no suppression of molting defects in intermediate loss-of-function *nekl* and *mlt* backgrounds that cause arrest at L2→L3 [*nekl-2(fd91), nekl-3(sv3)*, and *mlt-4(sv9)*] (Figure 5A), indicating that suppression by L2d may be allele specific.

**Figure 5.**
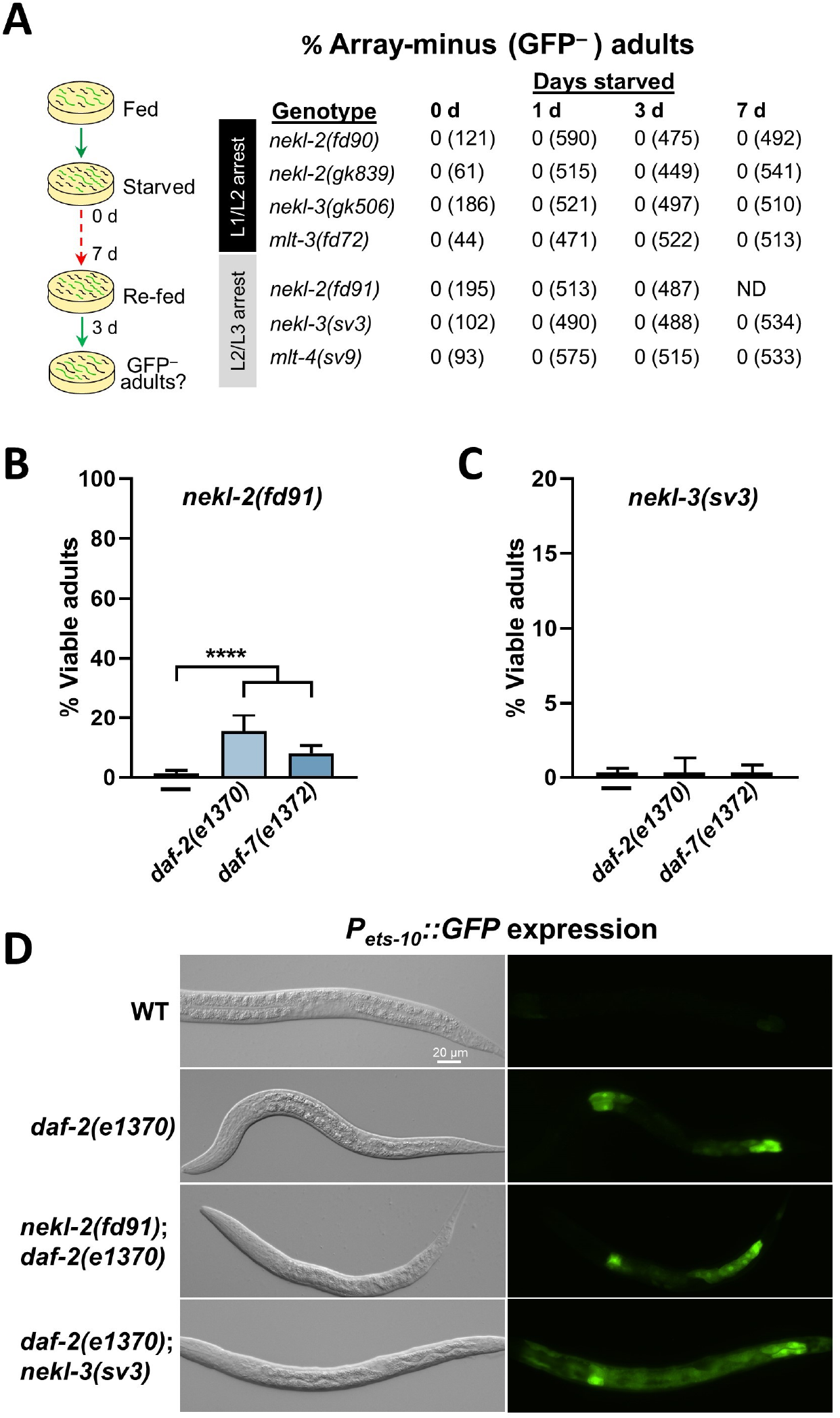
Transit through L2d does not suppress strong loss-of-function alleles of *nekl-2* or *nekl-3*. (A) Mixed-stage populations of null [*nekl-2(fd90), nekl-2(gk839), nekl-3(gk506)*, and *mlt-3(fd72)*] and intermediate loss-of-function [*nekl-2(fd91), nekl-3(sv3)*, and *mlt-4(sv9)*] mutants were starved for 0, 1, 3, and 7 days, and the percentage of viable array-minus (GFP^−^) adults was scored as described in Figure 2B. The total number of worms is shown in parentheses after each percentage (ND, not determined.) (B, C) Bar graphs showing the percentage of viable adults in *nekl-2* (B) and *nekl-3* (C) intermediate loss-of-function backgrounds containing the *daf-2* or *daf-7* Daf-c mutations. (D) Representative images of L2 or L2d larvae expressing the *P*_*ets-10*_*::GFP* reporter in the assayed backgrounds. Ten or more L2-stage worms were scored for the expression of the *P*_*ets-10*_*::GFP* reporter under a compound fluorescence microscope. Error bars represent 95% confidence intervals. Statistical significance was determined using Fisher’s exact test; ^****^p < 0.0001.

As a further test, we examined the ability of mutations in *daf-2* and *daf-7* to suppress L2→L3 molting defects in *nekl-2(fd91)* and *nekl-3(sv3)* intermediate loss-of-function mutants. Both *daf-2* and *daf-7* led to weak, but above background, suppression of *nekl-2(fd91*) but failed to suppress molting defects in *nekl-3(sv3)* mutants (Figures 5B–C), which may cause more severe molting defects than *fd91* (Lažetić & Fay, 2017a; Yochem et al., 2015). One potential explanation for the relative lack of suppression of *nekl-2(fd91)* and *nekl-3(sv3)* mutants by starvation or mutations is that they are defective at entry into L2d. To determine whether this is the case, we tracked the expression of *P*_*ets-10*_*::GFP* in *nekl-2(fd91)*; *daf-2(e1370)* and *daf-2(e1370)*; *nekl-3(sv3)* larvae. Notably, we observed L2d-like expression of *P*_*ets-10*_*::GFP* in 100% of second-stage larvae harboring the Daf-c mutations (Figure 5D). Collectively, our results indicate that the stage at which arrest occurs, although a factor, is not the only criterion required for L2d-mediated suppression. Rather, these data suggest that L2d can induce suppression when NEKL and MLT function has not been compromised beyond a critical threshold.

### Different molting stages have different requirements for NEKLs

The ability of *nekl-2*; *nekl-3* double mutants to transit the L2d→L3 molt but not the L2→L3 molt led us to hypothesize that the requirements for NEKL-2 and NEKL-3 may vary between different molts. To investigate the requirement for NEKL kinases at different larval-stage molts, we used the auxin-inducible degradation (AID) system to rapidly deplete CRISPR-tagged *nekl-2::aid* and *nekl-3::aid* strains at specific time points close to the start of each larval stage (Figure 6A) (Holland et al., 2012; Joseph et al., 2020; Zhang et al., 2015). In the case of NEKL-2, we observed substantial molting defects in all subsequent molts, indicating that NEKL-2 is required for all four molts during continuous development (Figures 6C). Unlike NEKL-2, depletion of NEKL-3 led to ∼100% molting defects only during the L1 L2 and L2 L3 molts, but only a small population of worms exhibited molting defects at later developmental stages (Figures 6D). This indicates that the functional requirement for NEKL-3 during molting is minimal after entry into the L3 stage. In contrast, treatment of wild-type control worms with auxin did not lead to growth arrest at any stage (Figure 6E). We also note that a substantial proportion of NEKL-3::AID worms underwent arrest at L1→L2 and L2→L3 molts in the absence of auxin treatment (Figure 6D). This is likely due to the inhibitory effects of the AID tag on NEKL-3 as well as some activity of the TIR1 ubiquitin ligase in the absence of auxin (Joseph et al., 2020; Lambrus et al., 2018; Natsume et al., 2016). Our data also suggest that neither NEKL-2 nor NEKL-3 is required for the L3d→L4 molt during the dauer exit process but that NEKL-2 is required for the post-dauer L4→adult molt (Figure 6C–D). As previously discussed, however, both NEKL-2 and NEKL-3 do appear to be necessary for the L2d→dauer molt. These findings indicate distinct NEKL requirements at specific molts, which may reflect different requirements for endocytosis at different larval stages.

**Figure 6.**
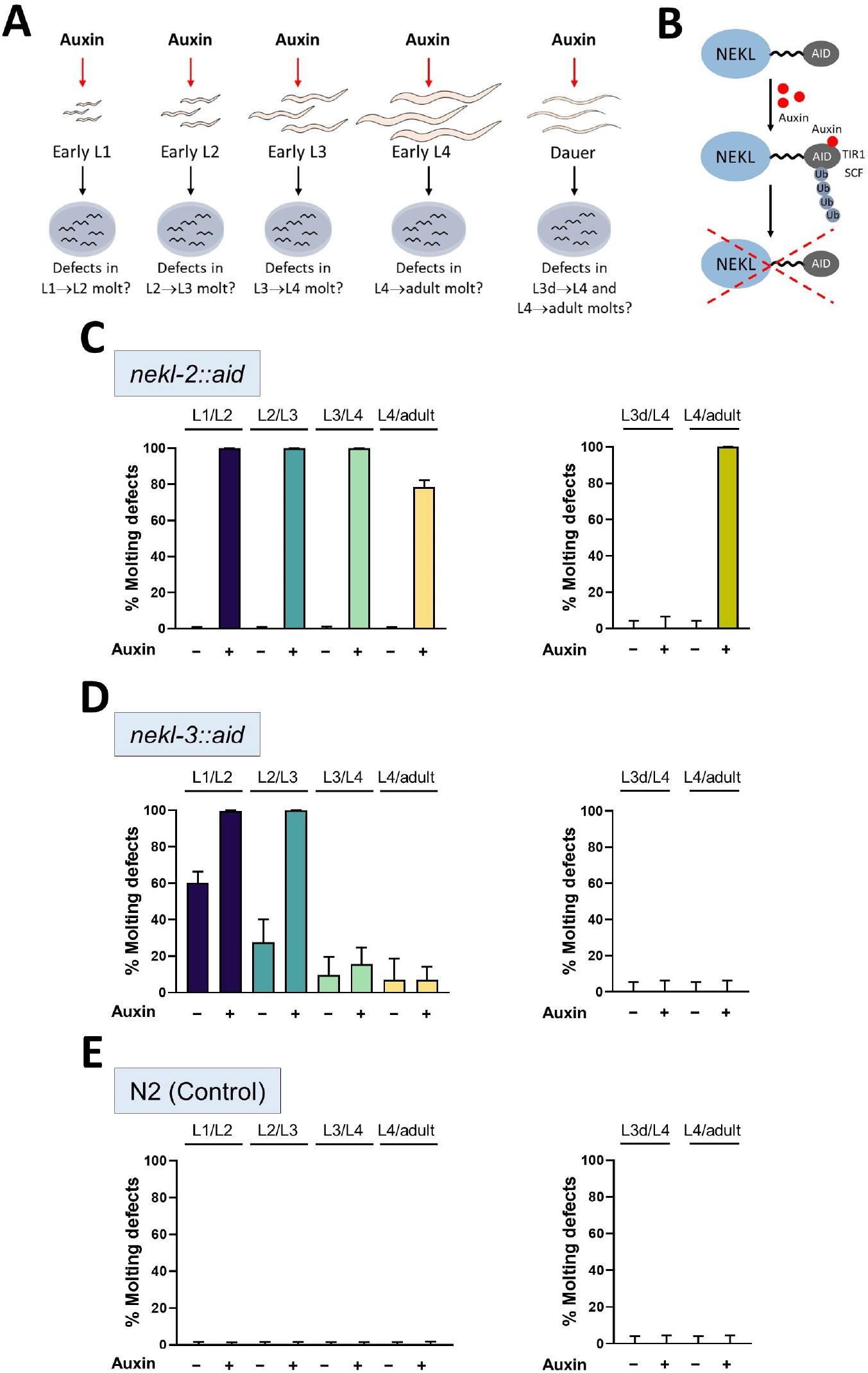
Requirements for NEKL kinases vary for different molts. (A) Schematic representation showing stages of larval development that were subjected to auxin-induced degradation of NEKLs (see Materials and Methods). (B) Schematic illustration showing how auxin-mediated degradation of NEKLs occurs. In the presence of auxin, TIR1-SCF complex binds to the AID tag and this interaction leads to ubiquitin-mediated degradation of NEKL::AID proteins. (C–E) Bar plots showing the percentage of molting-defective worms during continuous (left) and dauer-interrupted (right) development in the presence and absence of auxin in *nekl-2::aid* (C), *nekl-3::aid* (D), and wild-type (N2) (E) worms. Error bars represent 95% confidence intervals. For additional details see Materials and Methods.

### The dauer developmental trajectory is unlikely to mitigate trafficking defects in *nekl* mutants

We previously reported that NEKLs are required for epidermal membrane trafficking and that depletion of NEKLs leads to the reduced internalization of the epidermal cargo LRP-1 (Joseph et al., 2020). Moreover, we identified strong genetic suppressors of *nekl* molting defects that are directly involved in clathrin-mediated endocytosis including mutations affecting AP2 clathrin-adaptor subunits as well as the AP2 allosteric activator FCHO-1 (Joseph et al., 2020).

Importantly, these trafficking-related suppressors reduce both *nekl*-associated molting defects as well as defects in clathrin mobility and LRP-1 internalization (Joseph et al., 2020). To determine whether L2d affects *nekl* trafficking deficits, we examined the localization of LRP-1 on the apical surface of hyp7 in *daf-2(e1370)*; *nekl-3::aid* post-dauer adults. We found no evidence that passage through L2d and dauer can correct trafficking defects in NEKL-3 depleted adults, suggesting that suppression by L2d occurs through a distinct mechanism (Figure S3). This distinction between trafficking-related suppressors and L2d is consistent with our observation that whereas loss of AP2 and FCHO-1 activity can suppress strong loss-of-function mutations in *nekl-2* and *nekl-3* (Joseph et al., 2020), transit through L2d does not. We note that experimental constraints allowed us to cleanly follow LRP-1 localization in adult worms only, and, although unlikely, it remains possible that L2d may affect LRP-1 trafficking specifically at this stage.

### The length of the second larval stage cannot account for suppression by L2d

L2 diapause has a characteristic extended intermolt period as compared with the rapid development of an L2 larva after its first molt (Gems et al., 1998; Karp, 2018). We hypothesized that the longer intermolt period of an L2d larva may provide an extended window for *nekl-2*; *nekl-3* mutants to complete processes required for successful molting. To determine whether the length of the L2d larval stage is important for the suppression of molting defects, we tested five alleles of *daf-2—e1371, m577, e1368, m41*, and *e1370*—that have been extensively characterized and that differ in their phenotypic severity (Gems et al., 1998; Ruaud et al., 2011). Using a pharyngeal pumping assay to measure the characteristic pauses indicative of the lethargus phase of molting (Lažetić & Fay, 2017b), we observed that the length of the second larval stage varied considerably among worms containing different *daf-2* alleles (Figure 7A–B). Namely, the length of the L2 stage in alleles *e1371, m577*, and *e1368* was not substantially longer than that in wild type (10–12 h versus 9 h, respectively), whereas alleles *m41* and *e1370* took ∼2–3 times longer than wild type to complete the L2 stage (Figure 7A–B). Interestingly, all alleles except for *e1371* suppressed *nekl-2*; *nekl-3* mutants to a similar extent (Figure 7C), demonstrating that *nekl* suppression does not correlate with L2d length. These results indicate that the longer L2 inter-molt stage is unlikely to explain the observed suppression of molting defects by L2d.

**Figure 7.**
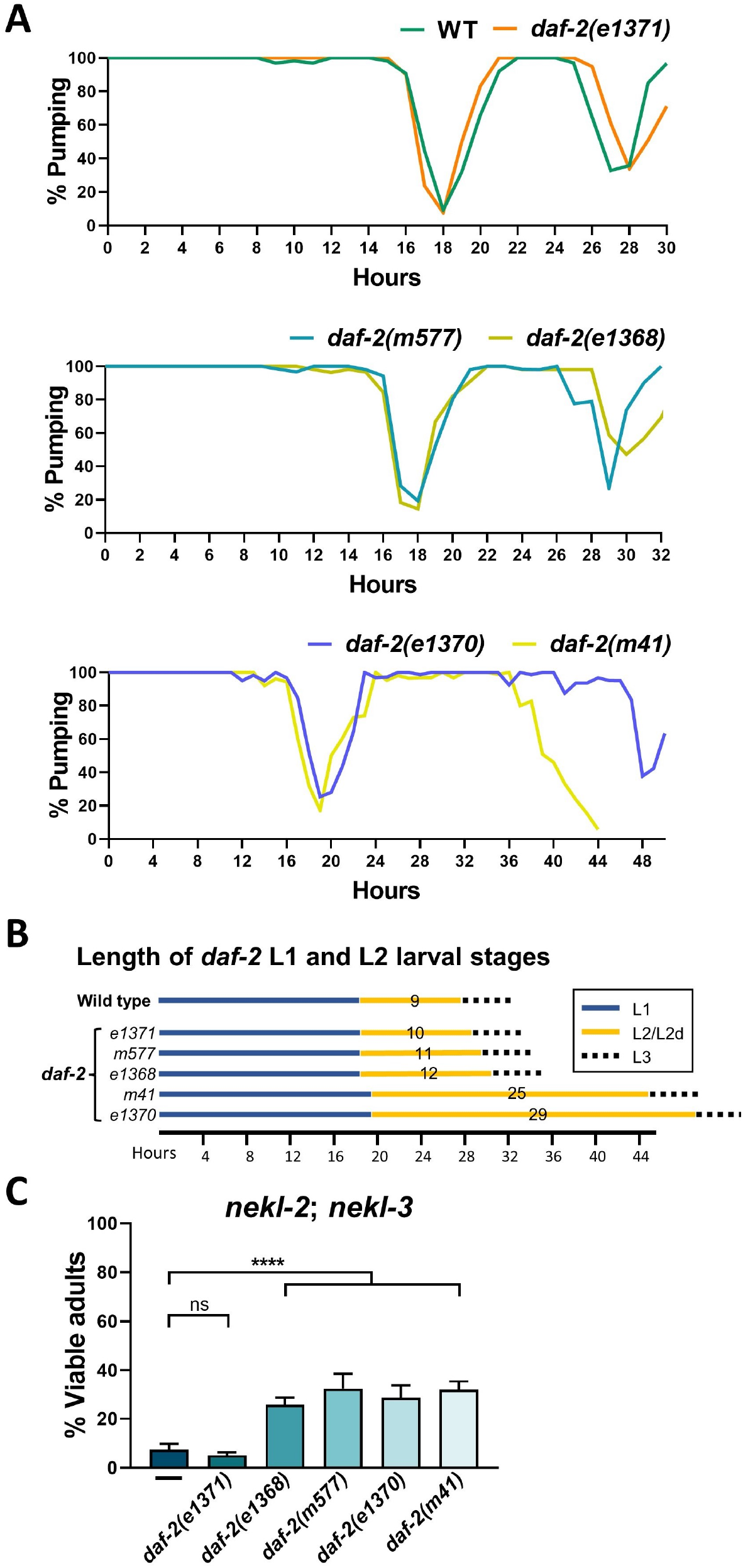
L2d length does not correlate with suppression by *daf-2* alleles. **A**. Pharyngeal pumping in *daf-2* alleles was used to determine the length of L1 and L2 larval stages. Experimental details are described in Materials and Methods. (B) Summary of data from panel A. (C) Bar graphs showing viable adults (%) scored in different *daf-2* mutant backgrounds. Error bars represent 95% confidence intervals. Statistical significance was determined by Fisher’s exact test; ^****^p < 0.0001. ns, not significant.

### NEKLs are required for the upregulation of cyclically expressed molting genes

Genetic studies have shown that steroid hormone signaling controls the activation of conserved nuclear hormone receptors, such as NHR-23 and NHR-25, which in turn regulate the expression of genes required for molting (Asahina et al., 2000; Kostrouchova et al., 2001; Kurzchalia & Ward, 2003; Lažetić & Fay, 2017b; Monsalve & Frand, 2012). Correspondingly, developmental time-course transcriptome analyses have demonstrated that thousands of genes in *C. elegans* oscillate during each larval stage and that the relative timing of their expression is consistent between molts (Hendriks et al., 2014; Meeuse et al., 2020; Turek & Bringmann, 2014). To better understand how NEKLs impact the molting process, we assessed the expression of four cyclically expressed genes (*mlt-10, nas-37, qua-1*, and *fbn-1*) that are required for molting in wild type and in *nekl* mutant backgrounds (Davis et al., 2004; Frand et al., 2005; Hao et al., 2006; Hendriks et al., 2014; Meeuse et al., 2020). To follow oscillating expression patterns, we used strains containing promoter fusions for each gene, with the promoter of interest fused to GFP tagged with the PEST sequence, which confers rapid degradation of the fluorescent marker (Frand et al., 2005; Hao et al., 2006; Meeuse et al., 2020).

As expected, wild-type synchronized larvae showed an oscillating pattern in accordance with previously published studies: P_*fbn-1*_::GFP-PEST and P_*mlt-10*_::GFP-PEST showed earliest and latest expression within each larval stage, respectively, with P_*nas-37*_::GFP-PEST expression occurring in the interval (Figure 8A–C and Figure S4A) (Hendriks et al., 2014; Meeuse et al., 2020; Turek & Bringmann, 2014). Notably, null mutations in *nekl-2* and *nekl-3* showed a dramatic reduction in the percentage of worms expressing these reporters during the L1 stage, as well as an ∼ 2-fold increase in the length of time before expression was first detected (Figure 8A–C and Figure S5– S6). For example, the first pulse of P_*mlt-10*_::GFP-PEST expression was detected in wild-type L1 larvae at ∼16 hours, whereas in null *nekl* backgrounds expression was detectable only after ∼ 32 hours and, unlike in wild-type larvae where it is expressed broadly, P_*mlt-10*_::GFP-PEST was confined to a small region in the anterior in mutants (Figure 8A and Figure S5). We also tested a fourth molting reporter, P_*qua-1*_::GFP::H2B-PEST, in null mutants and observed a substantial delay in L1 expression (Figure S4B–C).

**Figure 8.**
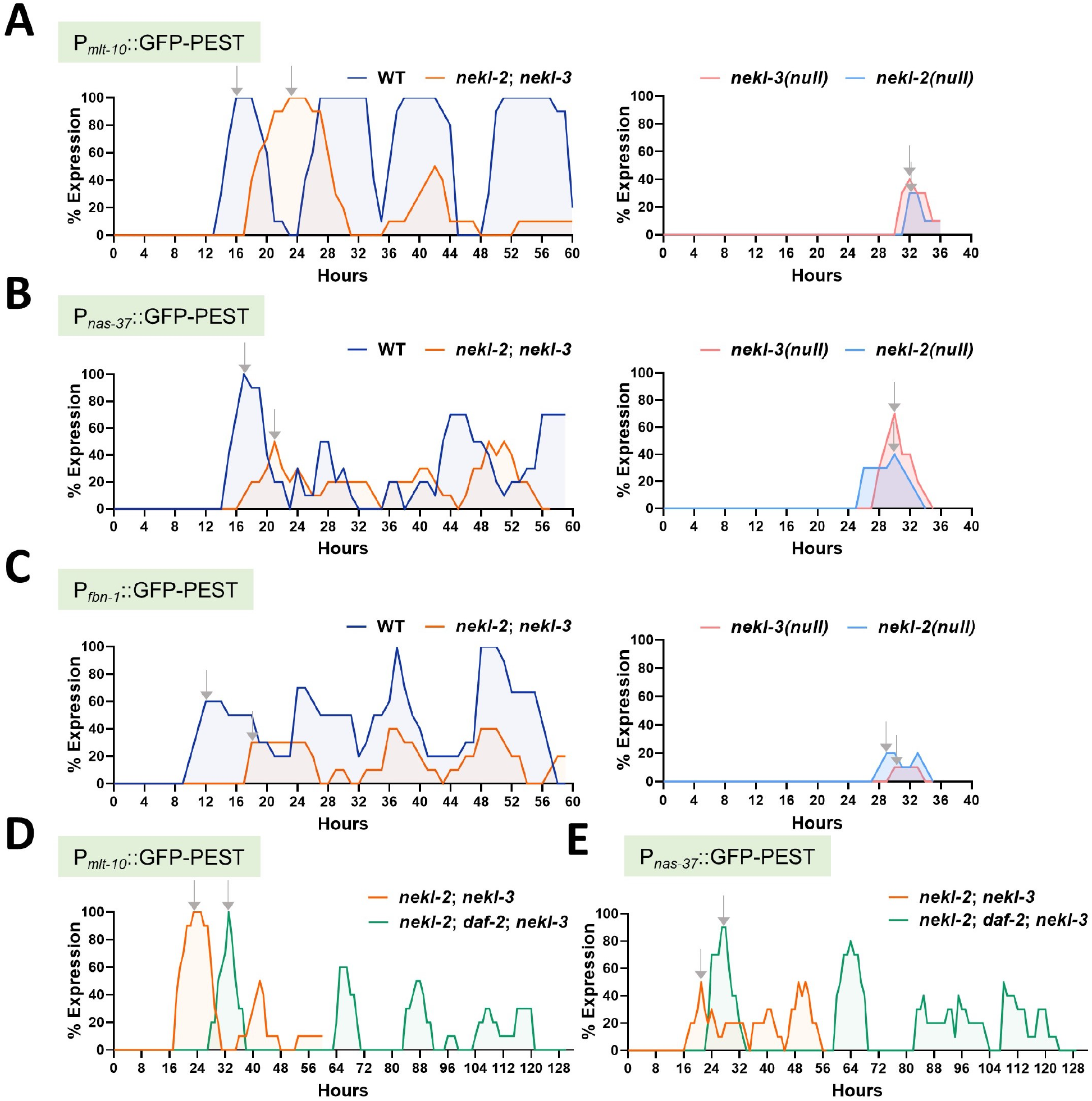
NEKLs are required for the synchronous and timely expression of cyclically expressed molting genes. (A–C) The percentage of worms expressing P_*mlt-10*_::GFP-PEST (A), P_*nas-37*_::GFP-PEST (B), and P_*fbn-1*_::GFP-PEST (C) was scored every hour (present or absent) after release from L1 arrest in synchronized cultures. Expression was observed in wild type, *nekl-2*; *nekl-3* double mutants, and null *nekl-2(gk839)* and null *nekl-3(gk506)* mutant worms using a high-powered dissecting microscope. (D,E) Expression of P_*mlt-10*_::GFP-PEST (D) and P_*nas-37*_::GFP-PEST (E) was followed in *nekl-2; nekl-3* mutants containing the *daf-2(e1370)* allele. In each experiment (A–E), at least 10 worms were scored every hour at 20°C for expression of the reporter. Grey arrow indicates peak expression of different reporters during late L1 stage.

In the case of *nekl-2*; *nekl-3* mutants, which arrest predominantly at the L2→L3 molt, we observed a slightly delayed but coherent peak of expression for P_*mlt-10*_::GFP-PEST during the L1 stage, followed by a less synchronous and delayed peak during the L2 stage (Figure 8A and Figure S5). We note that despite showing low/variable expression of P_*mlt-10*_::GFP-PEST for the remainder of the time course, these *nekl-2*; *nekl-3* animals remained arrested at the L2→L3 molt. Likewise, *nekl-2*; *nekl-3* strains containing P_*fbn-1*_::GFP-PEST and P_*nas-37*_::GFP-PEST markers also exhibited a slight delay in L1 expression but, more importantly, displayed highly variable and asynchronous expression of these markers at the L2 stage and in subsequent L2→L3 arrested larvae (Figure 8B–C). We note that although the time course of P_*fbn-1*_::GFP-PEST and P_*nas-37*_::GFP-PEST worms in *nekl-2*; *nekl-3* mutants appear to show some fluctuations in these markers, these worms remained arrested at the L2→L3 molt.

We also examined P_*mlt-10*_::GFP-PEST in *nekl-2(fd91)* and *nekl-3(sv3)* mutants, which also arrest at the L2→L3 molt, and observed a failure to upregulate P_*mlt-10*_::GFP-PEST specifically during the second larval stage (Figure S4D). Moreover, the ability of *nekl-2(fd91)* and *nekl-3(sv3)* to activate P_*mlt-10*_::GFP-PEST during L2 was reduced as compared with *nekl-2*; *nekl-3* strains, consistent with these alleles causing a more penetrant molting defect than the double mutant (Lažetić & Fay, 2017a; Yochem et al., 2015). Collectively, these observations demonstrate a strong correlation between the specific molting phenotypes of different *nekl* alleles and corresponding stage-specific defects in the expression of molting genes. Moreover, these results suggest that molting defects in *nekl* mutant backgrounds may be due to the reduced or delayed expression of genes required for the molting process.

Given the above observations, we were curious to know if the delays observed in the timing of the expression of molting genes in *nekl* mutants were specific to the molting process or whether other developmental processes might also be delayed in *nekl* mutants. To assess this, we made use of the Q cell system, which undergoes a stereotypical pattern of cell divisions, migrations, and differentiation into neurons during the L1 stage (Figure S7; Middelkoop & Korswagen, 2014). Based on previously defined stages, we scored Q cell developmental progression in wild-type, *nekl-2* null, and *nekl-3* null L1 larvae. Notably, we observed strong delays throughout Q cell development, with final differentiation into neurons taking ∼ 1.5-fold longer in *nekl* mutants than in wild type. These results suggest that reduced NEKL function not only delays the onset of molting gene expression but also slows the overall rate of development in these animals.

### *nekl-2*; *nekl-3* transcriptional defects are bypassed by progression through L2d

Our above results indicate that defects in the expression of molting genes during the second larval stage may account for the L2→L3 larval-arrest phenotype of *nekl-2*; *nekl-3* mutants. Given that L2d can suppress *nekl-2*; *nekl-3* molting defects, we were interested to determine whether transit through the L2d→L3 program also rescues expression defects in *nekl-2; nekl-3* mutants. Consistent with the suppression of molting defects, we observed clear and synchronous L1 and L2d expression of P_*mlt-10*_::GFP-PEST and P_*nas-37*_::GFP-PEST in *nekl-2*; *nekl-3* larvae that were shunted through L2d→L3 using a *daf-2* loss-of-function mutation (Figure 8D–E). Furthermore, these animals showed coherent expression peaks at the L3→L4 and L4→adult transitions, consistent with progression through the remainder of larval development. We note, however, that the *daf-2* mutation did not reverse delays in molting gene expression and, if anything, exacerbated this effect (Figure 8D–E). More broadly, we conclude that passage through L2d→L3 can sufficiently overcome transcriptional defects in *nekl-2*; *nekl-3* mutants, leading to the suppression of molting defects.

## Discussion

### Different requirements for NEKLs at different molts

In this study, we showed that developmental choices can affect molting outcomes in *C. elegans*. We found that starvation, exposure to dauer pheromone, and Daf-c mutations—conditions that induce L2d—all led to suppression of L2→L3 molting defects in *nekl-2*; *nekl-3* mutants. Starvation and dauer formation modulate other developmental processes in *C. elegans*, such as vulval induction (Battu et al., 2003; Baugh & Hu, 2020; Ferguson & Horvitz, 1985). In addition, passage through L2d and dauer corrects heterochronic defects (Abrahante et al., 2003; Ferguson & Horvitz, 1985; Liu & Ambros, 1991), in part by rewiring the gene regulatory network that controls temporal patterning (Ilbay & Ambros, 2019). In contrast, our findings suggest that the observed suppression of molting defects occurs because of different requirements for NEKL activities at different molts.

Figure 9A depicts a summary of our findings for NEKL-2 and NEKL-3 requirements at each molt. For example, timed-inactivation studies using NEKL-tagged AID strains revealed requirements for NEKL-2 and NEKL-3 at both the L1→L2 and L2→L3 molts, whereas only NEKL-2 is required for progression through the L3→L4 and L4→adult molts. In addition, the inability of partial loss-of-function alleles of *nekl*s to produce dauers under starvation conditions that induce L2d, indicates a requirement for both NEKL-2 and NEKL-3 in the L2d→dauer molt. Depleting NEKLs in dauer larvae and then moving those larvae to plates with bacteria also suggests that neither NEKL-2 nor NEKL-3 is required for the L3d→post-dauer L4→molt, although NEKL-2 is still required for the post-dauer L4→post-dauer adult molt.

**Figure 9.**
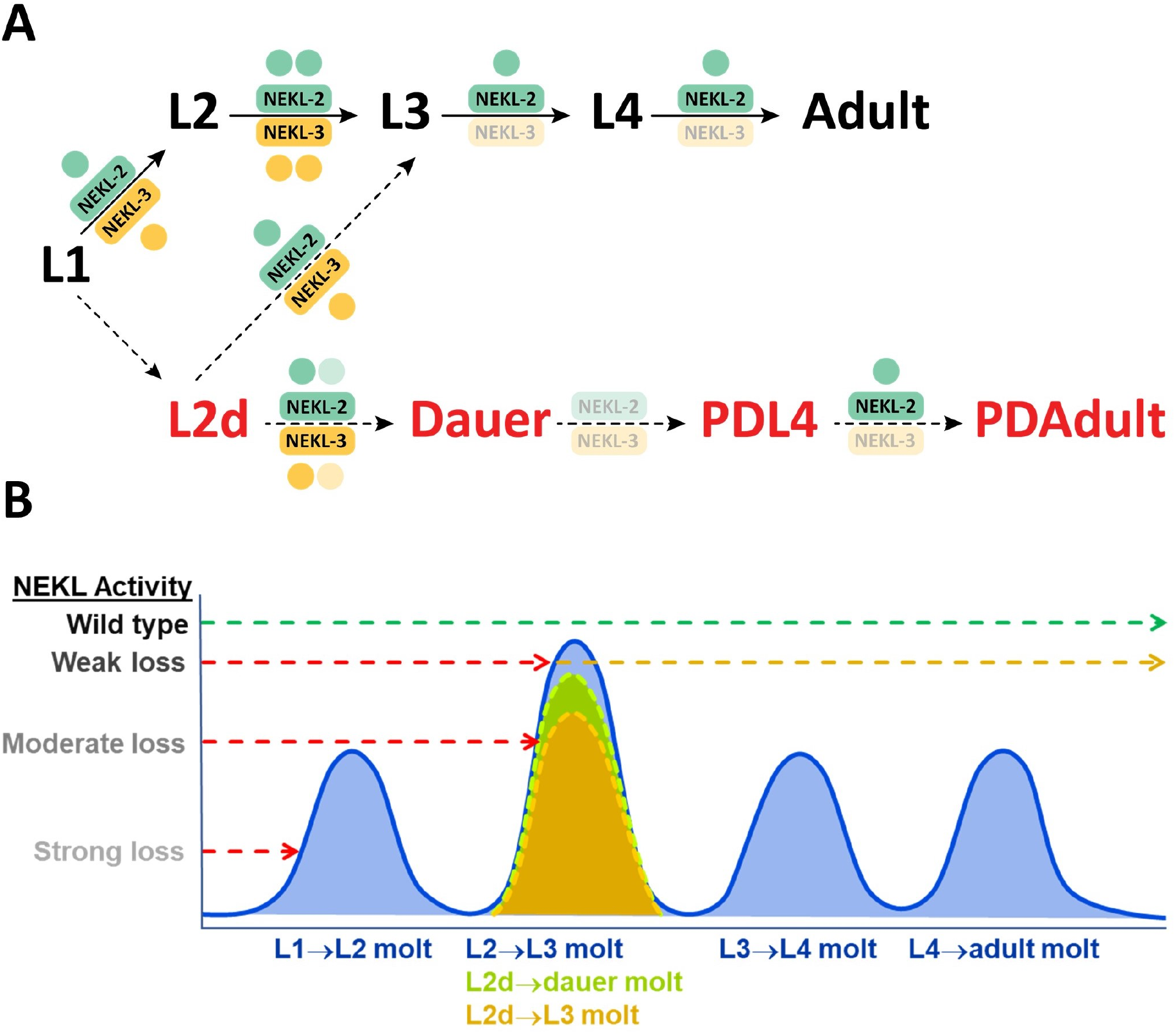
Summary of NEKL requirements at different molts. (A) Whereas NEKL-2 was required for all four larval molts, NEKL-3 was primarily required for the L1→L2 and L2→L3 molts. The requirement for NEKL kinases during the L2→L3 molt is higher (two dots) than during the L1→L2, L2d→L3 or L2d→dauer molts (fewer than two dots). Neither NEKL-2 nor NEKL-3 appears to be required for the molt immediately following dauer exit (dauer L4). The requirements for NEKL-2 and NEKL-3 during the final molt to adulthood appear similar for worms under continuous growth conditions or post dauer. PD, post-dauer. (B) A graphical representation showing threshold requirements for NEKL kinase activities at different molts. Whereas weak loss-of-function *nekl-2*; *nekl-3* mutants arrested at the L2→L3 molt, they were able to transit the L2d→L3 molt and, in the case of ascaroside treatment, the L2d→dauer molt, indicating that these latter two transitions have a lower threshold requirement for NEKL kinases. Strong reduction in kinase activity in intermediate loss-of-function mutants [*nekl-2(fd91)* and *nekl-3(sv3)*] was still sufficient to complete the L1→L2 molt, however, these mutants were unable to complete either L2→L3 or L2d→L3 molt. Finally, complete loss of kinase activity in null mutants [*nekl-2(gk839)* and *nekl-3(gk506)*] led to arrest at the L1→L2 molt.

In addition, genetic data which are summarized in Figure 9B, indicate that there is a greater requirement for NEKL-2 and NEKL-3 in the L2→L3 molt than in the L1→L2 molt. Specifically, *nekl-2*; *nekl-3* double mutants, along with *nekl-3(sv3)* and *nekl-2(fd91)* single mutants, exhibited molting defects at the L2→L3 molt, indicating that they were able to successfully transit the L1→L2 molt despite the partial reduction in NEKL activity. This observation also correlates with the expression of molting genes in these *nekl* mutant backgrounds, which showed a slightly delayed but nevertheless clear upregulation of molting genes during L1 but a failure to consistently upregulate these genes during L2. Lastly, the ability of L2d to suppress molting defects in *nekl-2*; *nekl-3* double mutants indicates that NEKL requirements are lower for the L2d→L3 transition than for L2→L3. Our studies also provide an explanation for how *nekl-2*; *nekl-3* mutants can successfully transverse the L3→L4 and L4→adult molts after completing L2d→L3. Namely, given that NEKL-3 is not required for the L3→L4 or L4→adult molts, the weak (aphenotypic) loss-of-function allele of *nekl-2, fd81*, would not be expected to cause arrest at these stages.

### The role of NEKLs in trafficking, transcription, and molting

By promoting the cyclic expression of hundreds of molting-related genes, steroid hormone signaling is a central feature of larval development. Likewise, the uptake of steroid hormone precursors, such as cholesterol, is essential for steroid hormone signaling. One such receptor, LRP-1, mediates the internalization of cholesterol by the epidermis and is required specifically in hyp7 for the completion of molting (Yochem et al., 1999). We previously showed that NEKL-2 and NEKL-3 regulate clathrin-mediated endocytosis and that depletion of NEKLs leads to the impaired endocytosis of LRP-1 at the apical membrane (Joseph et al., 2020). In the current study, we found that the expression of cyclically regulated molting genes is misregulated in *nekl* mutants and that these expression defects correspond closely with the strength of *nekl* alleles and with the timing of their developmental arrest points. Collectively, these findings imply that loss of NEKLs impairs steroid hormone signaling and thus the ability of larvae to transcribe genes required for molting.

In this study we found that transit through the uncommitted L2d stage (L2d→L3) can suppress both molting and transcriptional defects in *nekl-2*; *nekl-3* mutants, indicating that developmental plasticity and environmental context can affect this phenotype. Experiments suggest that L2d suppression is not due to either a correction of trafficking defects or an increase in the length of this larval stage. Rather, we infer that different stages of larval development may have distinct requirements for the level of steroid hormones and that the threshold for certain transitions, such as L1→L2 and L2d→L3, may be less than that for other transitions, such as L2→L3. Alternatively, it is possible that during certain life stages a sufficient level of steroid-hormone building blocks is naturally present, such that there is less dependence on importing additional signaling precursors via NEKLs and epidermal endocytosis. Consistent with this possibility, neither NEKL-2 nor NEKL-3 was required for the dauer→L4 transition, and dauer larvae are known to concentrate lipids (Jeong et al., 2005; Popham & Webster, 1979).

Future studies will seek to understand precisely how NEKL kinases regulate targets involved in intracellular trafficking and will increase our knowledge of which epidermal trafficking cargoes are critical for molting.

## Acknowledgments

We thank Amy Fluet for editing this manuscript and Eric Jorgenson, Paul Sternberg, Alison Frand, and Helge Grosshans for strains. This project was supported by R35 GM136236 to DSF and by an Institutional Development Award (IDeA) from the National Institute of General Medical Sciences of the National Institutes of Health (P20GM103432).

