## Supplementary Figures 1-7 for "A life cycle alteration can correct molting defects in *Caenorhabditis elegans*"

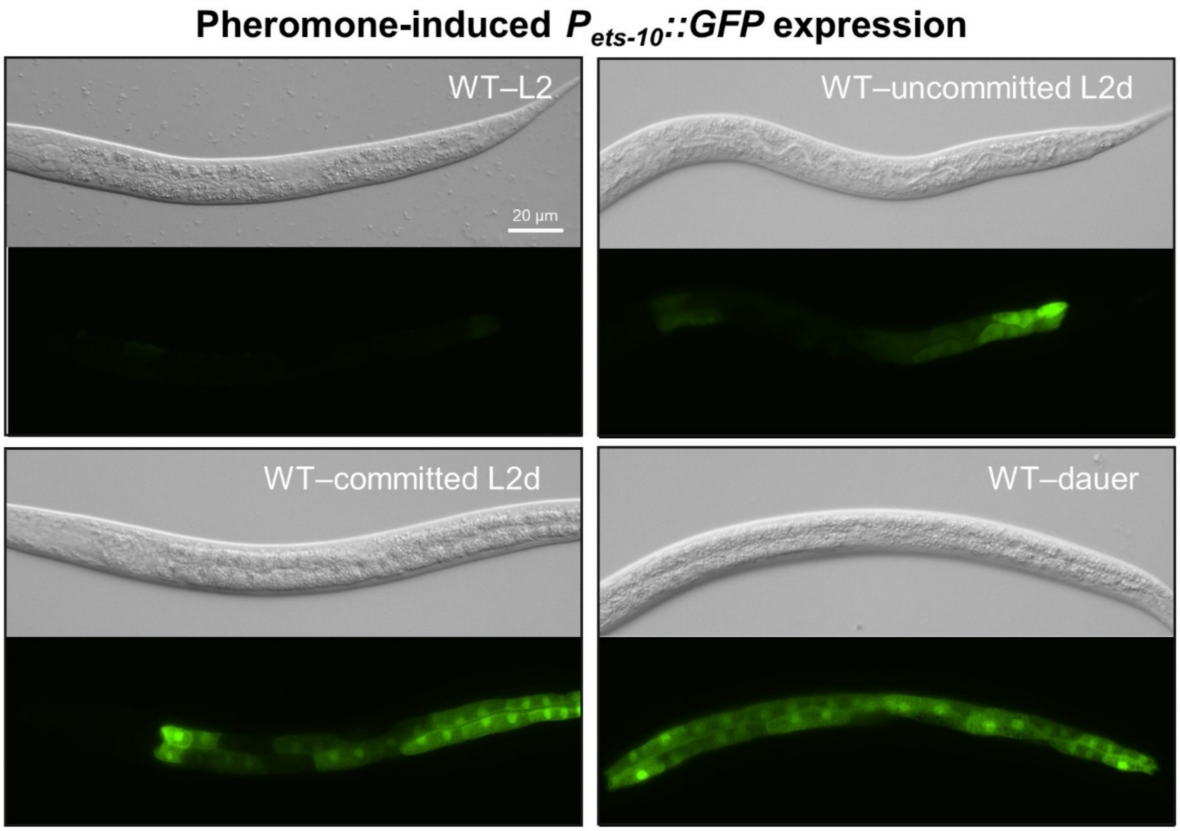


**Figure S1. Expression of *$P_{ets\text{-}10}$::GFP* in the presence of ascarosides.** Representative images of wild-type (WT) L2 (no ascarosides), uncommitted L2d, committed L2d, and dauer animals expressing *$P_{ets\text{-}10}$::GFP*. Images were taken using a 40× compound scope objective.

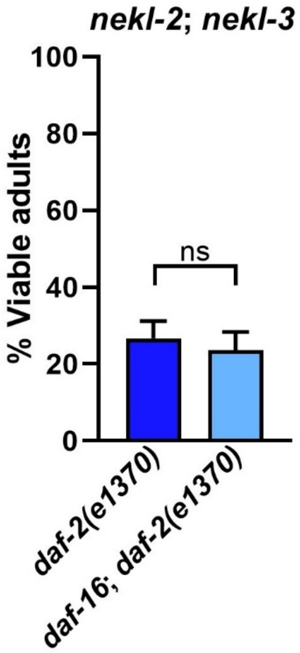


**Figure S2. Suppression of molting defects by *daf-2* was not significantly inhibited by *daf-16*.** Bar graph showing the viable adults (%) when the indicated mutants were grown at 20°C. Error bars represent 95% confidence intervals. ns, not significant.

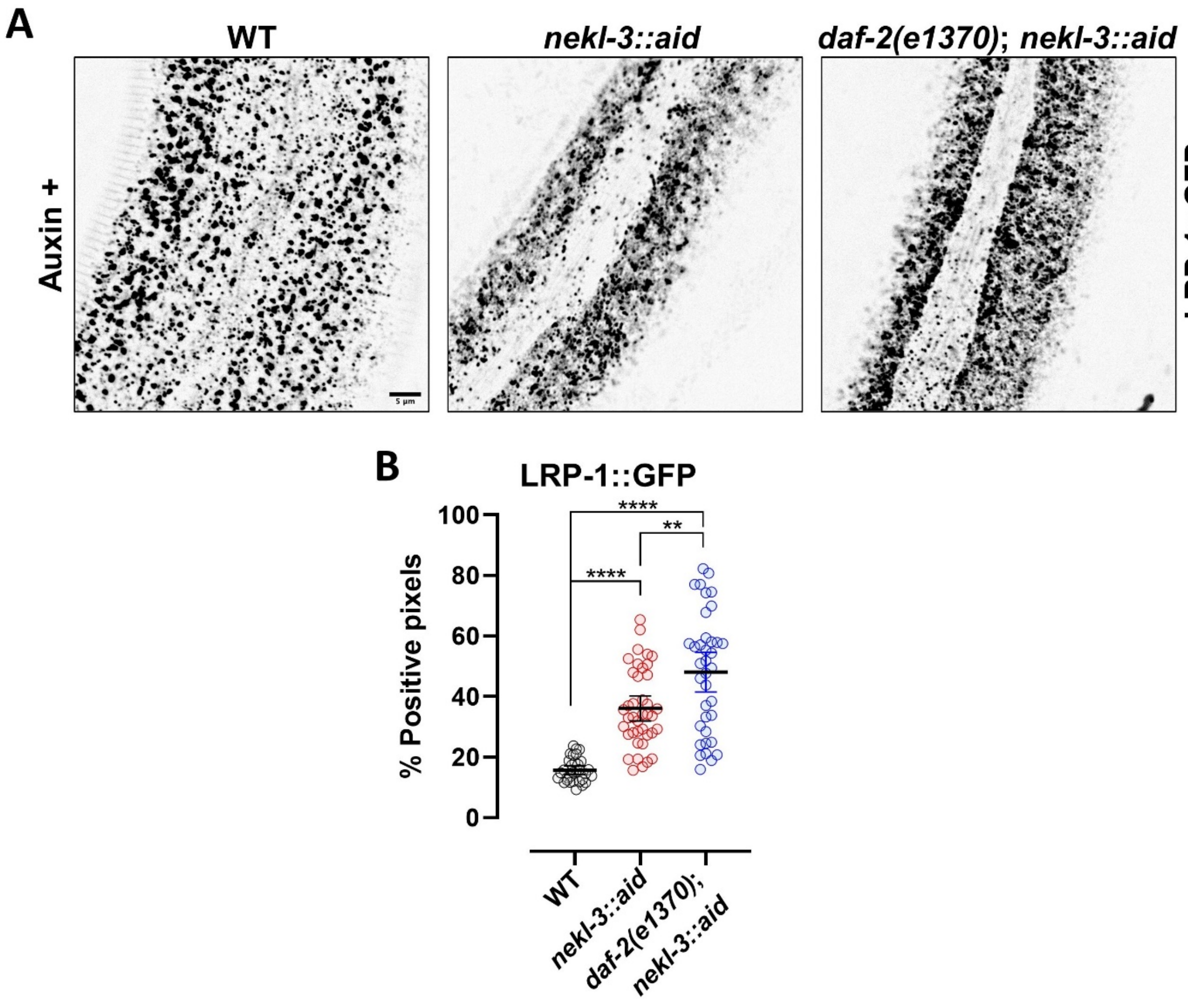


**Figure S3. Localization of LRP-1 in NEKL-3–depleted adults.** (A) Representative images of auxin-treated day 2 adults expressing LRP-1::GFP in the apical region of hyp7 in the indicated mutant backgrounds. (B) Percentage of GFP-positive pixels above threshold for each adult and the means for each background are plotted. Error bars represent 95% confidence intervals. Statistical significance was determined using a two-tailed, unpaired t-test; $**p < 0.01$, $****p < 0.0001$.

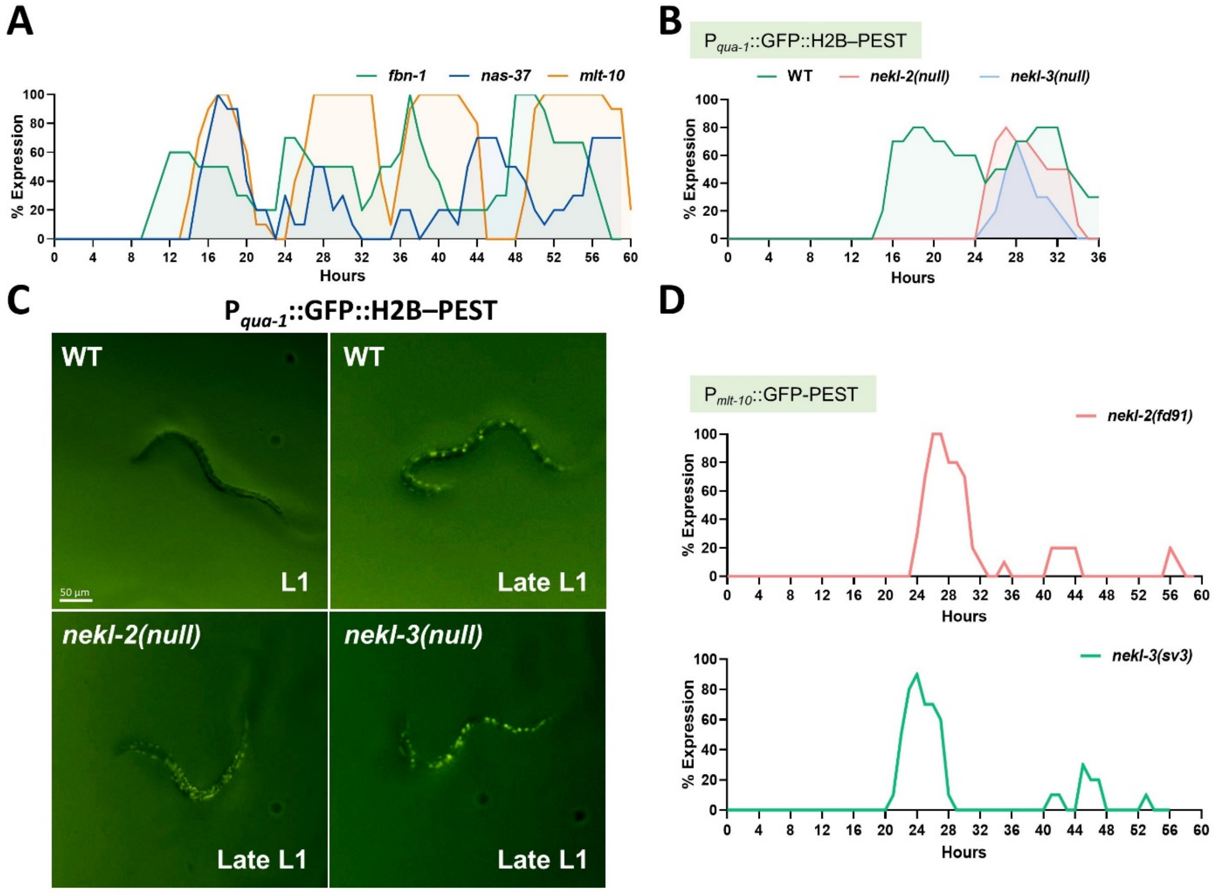


**Figure S4. Expression of molting genes in different backgrounds.** (A) Expression of $P_{fbn-1}$::GFP-PEST *,* $P_{nas-37}$::GFP-PEST, and $P_{mlt-10}$::GFP-PEST in wild type showing their initial peak expression at different phases. (B) Expression of $P_{qua-1}$::GFP::H2B–PEST was followed for 36 hours in wild type and null *nekl-2(gk839)* and *nekl-3(gk506)* mutants. (C) Representative images showing wild type and null *nekl-2 (gk839)* and *nekl-3(gk506)* OR *nekl-2 (null)* and *nekl-3(null)* mutants expressing $P_{qua-1}$::GFP::H2B–PEST before the L1 molt. (D) Expression of $P_{mlt-10}$::GFP-PEST was followed in *nekl-2(fd91)* and *nekl-3(sv3)* mutants, which arrest at the L2→L3 molt. For experiments in panels A, B, and D, at least 10 worms were scored every hour for expression of the indicated reporters (present or absent) following release from L1 arrest.

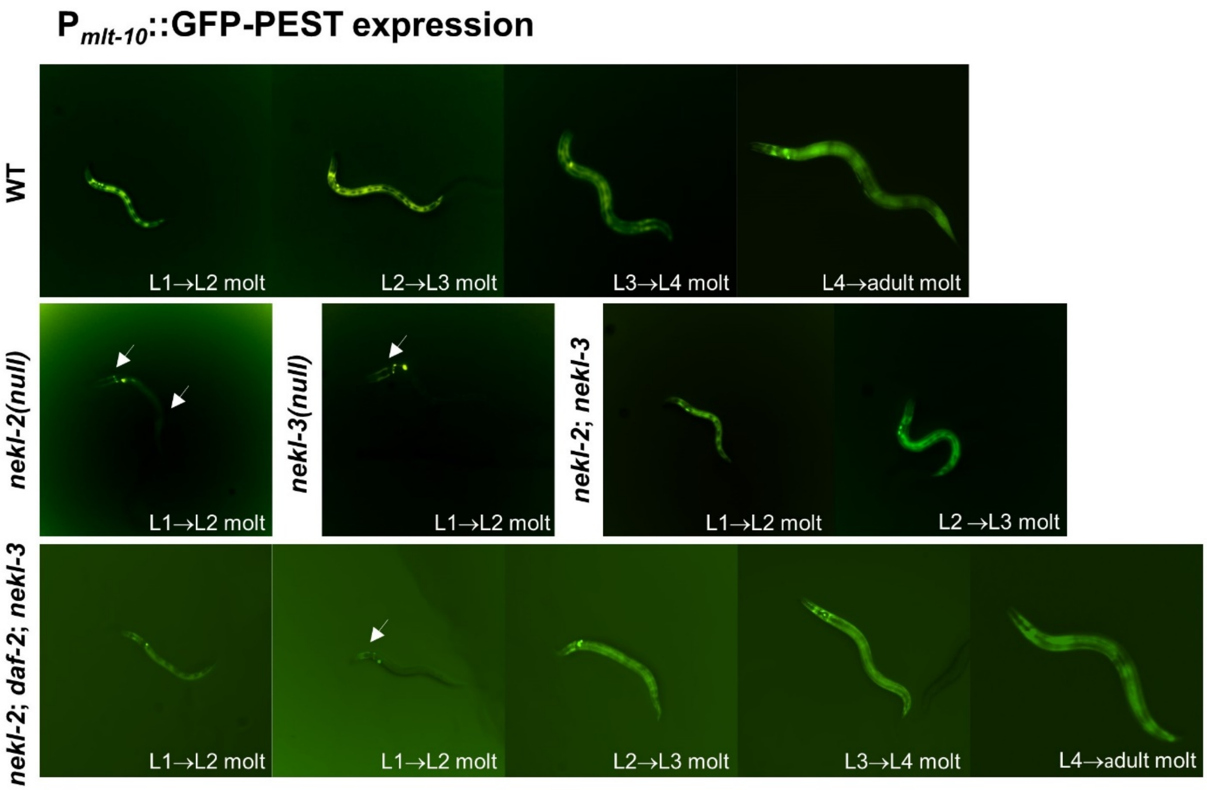


**Figure S5. Differential expression of P$_{mlt-10}$::GFP-PEST in different *nekl* mutants**. Representative images of larvae expressing P$_{mlt-10}$::GFP-PEST in wild type, null *nekl-2(gk839)*, null *nekl-3(gk506)*, *nekl-2*; *nekl-3*, and *nekl-2*; *daf-2*; *nekl-3* backgrounds during different stages of development at 20°C. White arrows indicate reduced/inconsistent expression of P$_{mlt-10}$::GFP-PEST in mutants as compared with wild type.

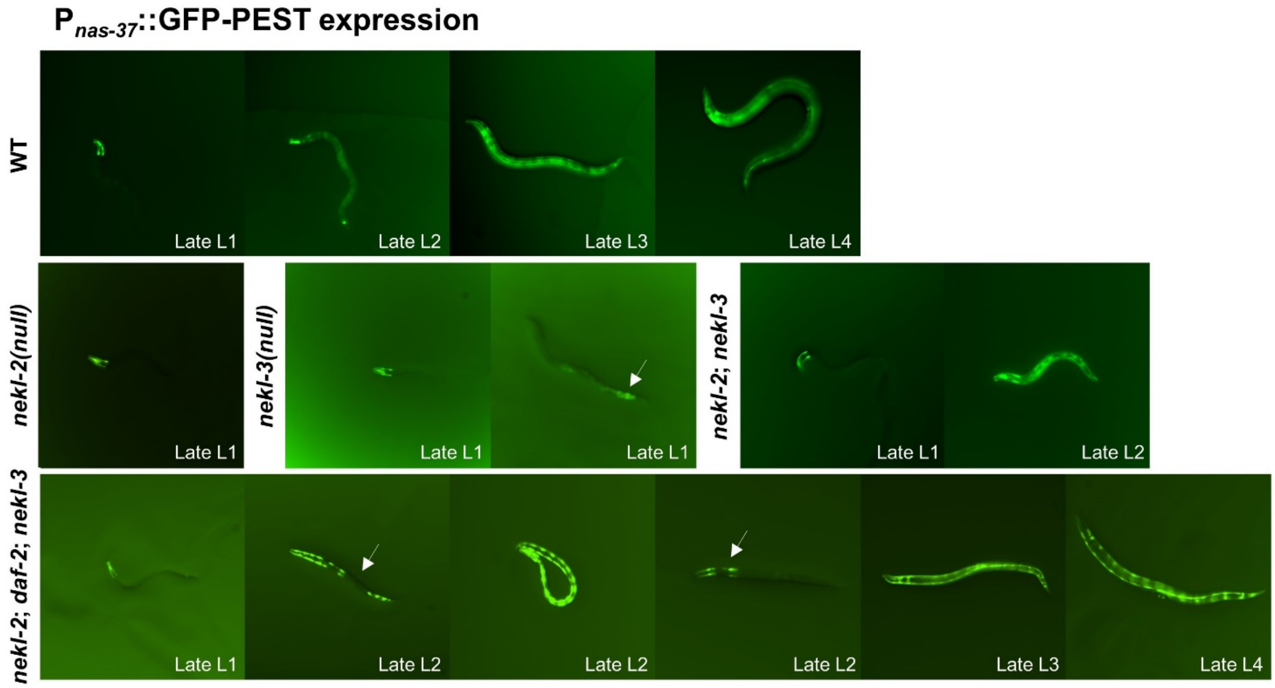


**Figure S6. Expression of P$_{nas\text{-}37}$::GFP-PEST in different *nekl* mutants**. Representative images of larvae expressing P$_{nas\text{-}37}$::GFP-PEST in wild type, null *nekl-2(gk839)*, null *nekl-3(gk506)*, *nekl-2; nekl-3*, and *nekl-2*; *daf-2*; *nekl-3* backgrounds at 20°C. White arrow indicates inconsistent expression of P$_{nas\text{-}37}$::GFP-PEST in mutants when compared with wild type. In the wild-type background, seam cell expression of P$_{nas\text{-}37}$::GFP-PEST was observed only in L4 larvae. Interestingly, the suppressed strain (*nekl-2*; *daf-2*; *nekl-3*) expressed P$_{nas\text{-}37}$::GFP-PEST in seam cells of L2, L3, and L4 larvae. This change in the expression pattern of P$_{nas\text{-}37}$::GFP-PEST could be an effect of the *daf-2* mutation.

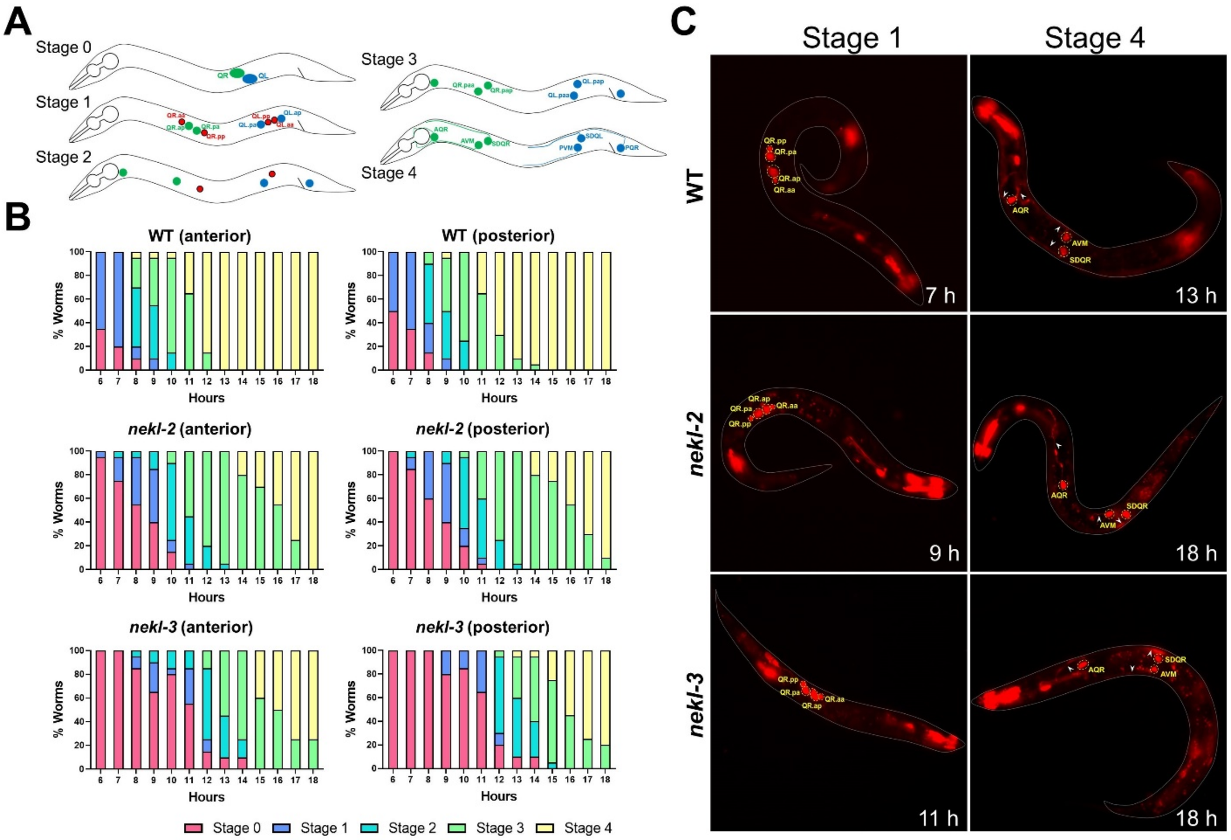


**Figure S7. Migration and differentiation of Q cell neuroblasts.** (A) Schematic showing the five stages of Q neuroblast development. Stage 0: Q neuroblasts (QR and QL) have not yet undergone divisions. Stage 1: Each Q cell has divided two times to generate two apoptotic cells, which are later engulfed by hyp7, and two neural progenitors (QR.ap, QR.pa, QL.ap, QL.pa). Stage 2: QR.ap migrates toward the pharynx, and QL.ap migrates toward the rectum. Stage 3: QR.pa and QL.pa each divide into two cells. Stage 4: The six remaining cells differentiate into neurons. (B) The stages of development of Q cells were scored every hour in wild type, *nekl-2(gk839)*, and *nekl-3(gk506)* mutants expressing $P_{egl\text{-}17}$::Myri-mCherry fusion protein, which is expressed in Q cells. Note that all stages are delayed in *nekl* mutants. In each experiment, 20 worms were scored every hour after release from L1 arrest. (C) Representative images of wild-type, *nekl-2*, and *nekl-3* worms expressing $P_{egl\text{-}17}$::Myri-mCherry at stage 1 (left column) and stage 4 (right column) of Q neuroblast development. The relevant cell types are labeled. Note that $P_{egl\text{-}17}$::Myri-mCherry is also expressed in some additional (unlabeled) cell types. White arrowheads show the differentiated neurons during stage 4 of Q cell development.
